# Population based selection shapes the T cell receptor repertoire during thymic development

**DOI:** 10.1101/2022.02.14.480309

**Authors:** Francesco Camaglia, Arie Ryvkin, Erez Greenstein, Shlomit Reich-Zeliger, Benny Chain, Thierry Mora, Aleksandra M. Walczak, Nir Friedman

## Abstract

One of the feats of adaptive immunity is its ability to recognize foreign pathogens while sparing the self. During maturation in the thymus, T cells are selected through the binding properties of their antigen-specific T-cell receptor (TCR), through the elimination of both weakly (positive selection) and strongly (negative selection) self-reactive receptors. However, the impact of thymic selection on the TCR repertoire is poorly understood. Here, we use transgenic Nur77-mice expressing a T-cell activation reporter to study the repertoires of thymic T cells at various stages of their development, including cells that do not pass selection. We combine high-throughput repertoire sequencing with statistical inference techniques to charactarize the selection of the TCR in these distinct subsets. We find small but significant differences in the TCR repertoire parameters between the maturation stages, which recapitulate known differentiation pathways leading to the CD4+ and CD8+ subtypes. These differences can be simulated by simple models of selection acting linearly on the sequence features. We find no evidence of specific sequences or sequence motifs or features that are suppressed by negative selection. These results are consistent with a collective or statistical model for T-cell specificity, where negative selection biases the repertoire away from self recognition, rather than ensuring lack of self-reactivity at the single-cell level.

## I. INTRODUCTION

In order to protect themselves against infection, jawed vertebrates have evolved an adaptive immune system. T lymphocytes play a leading role in this system. Each T lymphocyte expresses a unique T-cell receptor (TCR) capable of binding short protein fragments presented by the host’s Major Histocompatibility Complexes (MHC), subsequently triggering clonal expansion and differentiation of immune effector function. The T cell system discriminates pathogen derived “foreign” proteins from the body’s own “self” proteins, in such a way that an immune response is usually triggered only by peptides from exposure to a potentially harmful threat. We ask if we can identify specific TCR features which allow the system to discriminate foreign and self-peptides.

TCRs are generated in a stochastic assembly process based on random recombinations of genomic templates and additional non-templated insertions and deletions [1]. The ability to discriminate between self and non-self targets cannot therefore be inherited, but must be learned afresh in each individual. This process is widely believed to occur during the development of haemopoetic precursors into mature T cells, which occurs in a specialized microenivironment within the thymus. This process has been studied in considerable detail. T cells precursors first produce a *β* chain and if the generated chain is functional, the cell proliferates and an *α* chain is generated. While the TCR chains are being assembled, CD4 and CD8 surface markers are expressed as precursor cells transit to the Double Positive state (DP). DP TCR are subject to thymic selection, a process that tests receptor binding by presenting them with the organism’s own proteins, and eliminates very weak binders (positive selection), but also too strongly self-reactive receptors (negative selection) [2]. During thymic selection, DP cells differentiate into CD4+ or CD8+ cells by keeping expression of only one of these molecules, which determines their function. While this picture is well-established and the maturation trajectory has a well established gene expression signature [3], the TCR sequences removed during thymic selection, which should be manifested as “holes” in the repertoire, have never been directly observed. The lack of quantifiable signatures of thymic selection, differentiation and proliferation hinders a dynamic description of TCR maturation [4].

Positive and negative selection imposes upper and lower boundaries on the binding energy of the interaction between TCR and self peptide-MHC complexes [5]. However, given the timescales of thymic selection [2], it is difficult to understand how a given TCR can be exposed to *all* possible self-peptides. It seems more likely that each cell probes only a random subset of self-peptides, which will vary for different T cells. Previous studies have estimated that each TCR probes ~ 1-5% of self-peptides in humans [6, 7]. Individual T cells may therefore experience their own idiosynchratic selection process, and remain cross-reactive to an overwhelming majority of self-peptides. Consistent with this model negative selection is known to be leaky, letting auto-reactive cells differentiate into regulatory T-cells [8, 9]. The partial nature of negative selection may limit its impact on the repertoire. However, the question of how systematic TCR elimination is during negative selection remains open.

The difficulty of characterizing negatively selected sequences is partly due to survivor bias—eliminated sequences cannot be seen when sampling functional immune repertoires [10–13]. To overcome this limitation, we sequenced the TCR repertoire of thymocyte subpopulations isolated from mice carrying a reporter transgene linked to Nur77, a marker of T cell activation both within the thymus and in the periphery. Nur77 expression, in combination with a marker of cell death, allows us to identify cells which do not pass thymic selection, as well as those that do, providing us with a window into the unselected repertoire. By comparing the sequenced repertoires to statistical models of mouse TCR generation [13], and subset-specific models of thymic selection [14], we searched for specific TCR sequence features that lead to positive and negative selection, at various stages of intra-thymic T cell development.

## II. RESULTS

### Tracking T cell development stages by flow cytometry

To identify specific sequence features of TCR during each step of thymic selection, we performed high-throughput sequencing of TCR repertoires from different subpopulations of thymocytes from transgenic Nur77 reporter expressing mice. These mice carry a fluorescent reporter gene which is co-expressed with Nur77, a marker of T cell activation. [15]. Three genetically identical Nur77 reporter mice were sacrificed at the age of 6 weeks, when thymus development is completed and its cell population is stable [16]. Thymus and spleen were removed, and stained for fluorescence-activated cell sorting (see Materials and Methods). The cells were sorted based on Nur77 reporter expression (to detect activation), Annexin V (to detect early apoptosis) in combination with CD3, CD4, and CD8 cell surface markers. We used the gating strategy illustrated in Figs. 1A, B, C to isolate double positive DP cells preceding selection (CD4+CD8+, Nur77-, Annexin V-: DP pre), DP cells in the process of being positively selected (CD4+CD8+, Nur77+ Annexin V-: DP pos), DP cells dying by neglect (CD4+CD8+, Nur77-Annexin V+: DP dbn); and single positive (SP) cells: CD4+CD8-, Nur77-Annexin V+ (CD4 negS), and CD4-CD8+, Nur77-Annexin V+ cells (CD8 negS), assumed to be undergoing negative selection. In addition, we sequenced the repertoires of mature (post-selection) single positive SP CD4+ and CD8+ cells from the spleen (CD4 spl and CD8 spl). The proposed differentiation pathway between these populations at different maturation stages are schematically represented in Fig. 1D. Together, these seven repertoires should contain both the selected thymocytes and the thymocytes which fail either positive or negative selection and die in the thymus.

**FIG. 1:**
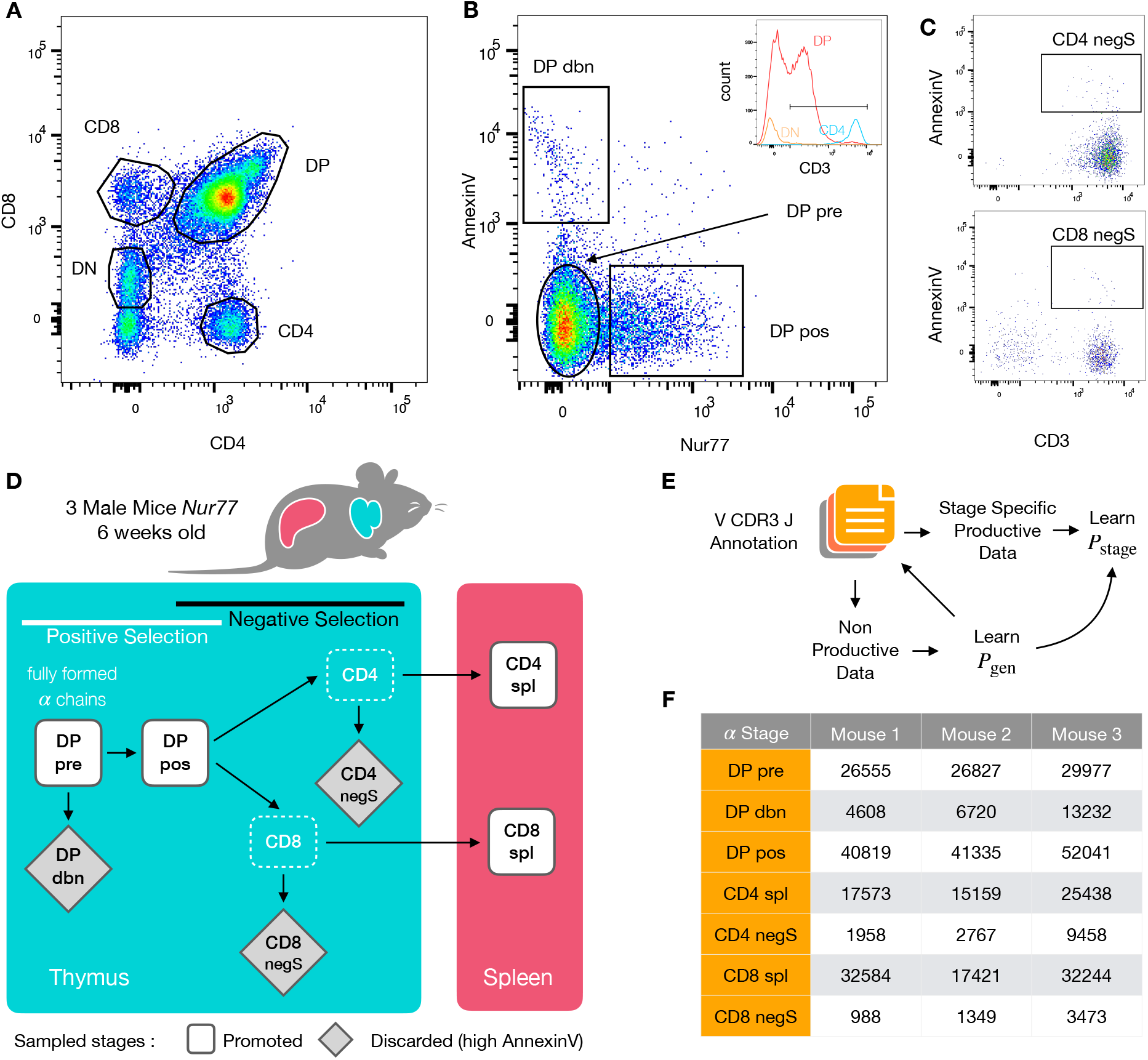
Experiment outline and repertoire sampling. **A** Flow cytometry scatterplots of T cell population from the thymus according to the markers CD4 and CD8. **B** The DP population is separated from DN according to CD3 expression (insert). Cells are then FACS sorted according to the expression of Nur77 and AnnexinV. **C** CD4 cells in the spleen (above) and CD8 (below) are FACS sorted according to the expression of CD3 and AnnexinV. **D** Schematic evolution of the sampled cell types during thymic maturation. **E** Analysis workflow: annotated reads in sampled repertoires are input for model inference (see Materials and Methods). Out-of-frame TCR sequences are pooled from all mice and stages to learn a generation model. Inframe sequences are used to learn maturation stage specific selection models with the generation model as background. **F** Numbers of unique in-frame *α* chain sequences obtained for the maturation stages in each mouse after annotation (see Fig. S6A for *β* chain).

### TCR repertoire sequencing

We sequenced and annotated the TCR repertoires of each subset as described in Materials and Methods. The cDNA of individual *α* and *β* genes (TRA and TRB) were barcoded with unique molecular identifiers (UMI) in order to allow for correction of sequencing errors and PCR bias. However, in this analysis we focused on unique sequences (discarding count information) to avoid expression and amplification biases. As a quality control of the whole procedure, we showed that the number of alpha and beta sequences within each population was highly correlated (Fig. S1). We further verified that the relative fraction of TRA sequences associated with iNKT cells (identified by TRAV11 and TRAJ18 genes [17]) is higher in CD4 than in CD8 cells (see Fig. S2).

We obtained seven datasets for both chains and for each of the 3 mice. A small fraction of sequences contain stop codons, usually because of a frameshift in the CDR3. These sequences likely come from transcription from a chromosome carrying a nonproductive chain, which is known to persist despite allelic exclusion acting on the TCR locus. The rest of the sequences are assumed to be productive. Since nonproductive TCR owe their survival to the productive gene on the other chromosome, they are not affected by selection. We combined all nonproductive sequences from all subsets to infer a generative mechanistic model of the V(D)J recombination process using IGoR [18]. Once trained, the model can be used to assign a generation probability *P*_gen_ to any TCR sequence observed [18, 19] (see Materials and Methods and Fig. 1E).

The datasets contain ~ 1,000 – 50,000 unique productive sequences per subset (Figs. 1F and S6A). Since the 3 mice were isogenic and shared the same MHC haplotype, we expect their repertoires to be subject to the same processes of recombination and selection [10]. Unless specified otherwise, all downstream analyses were therefore carried out on pooled productive TCR sequences from each population from the three individuals to increase statistical power.

### Repertoires from different T cell populations have different statistical parameters

To assess how selection acts at the different maturation stages, we studied the distribution of sequence features in TRA repertoires. We compared TRAV and TRAJ gene usage at the different maturation stages with each other and with their excepted frequency from the generation model learned from nonproductive TCR sequences, which we will refer to as the pre-selection model or *P*_gen_ (Figs. 2A, S3). TRAV usage broadly follows the pattern of the pre-selection model, although SP CD4+ repertoires have a lower proportion of TRAV12-2, and most populations have an increased proportion of TRAV7-2. TRAJ gene usage also broadly agrees with the pre-selection model predictions, although SP CD8+ repertoires have a lower proportion of TRAJ31, SP CD4+ repertoires have an increased proportion of TRAJ27 and TRAJ32 which is underrepresented in all cell types (Fig. S3). For both V and J genes, we see little difference between the repertoires of spleen CD4 and CD8 cells, and their discarded counterparts in the thymus (negS). We also observe strong similarities between all the DP subsets. TRB gene usage follows similar trends, although there are some differences in J gene usage between selected and unselected SP CD4+ and CD8+ cells. Overall the biases of the recombination process dominate any effects of selection on V and J region usage. (Fig. S5).

**FIG. 2:**
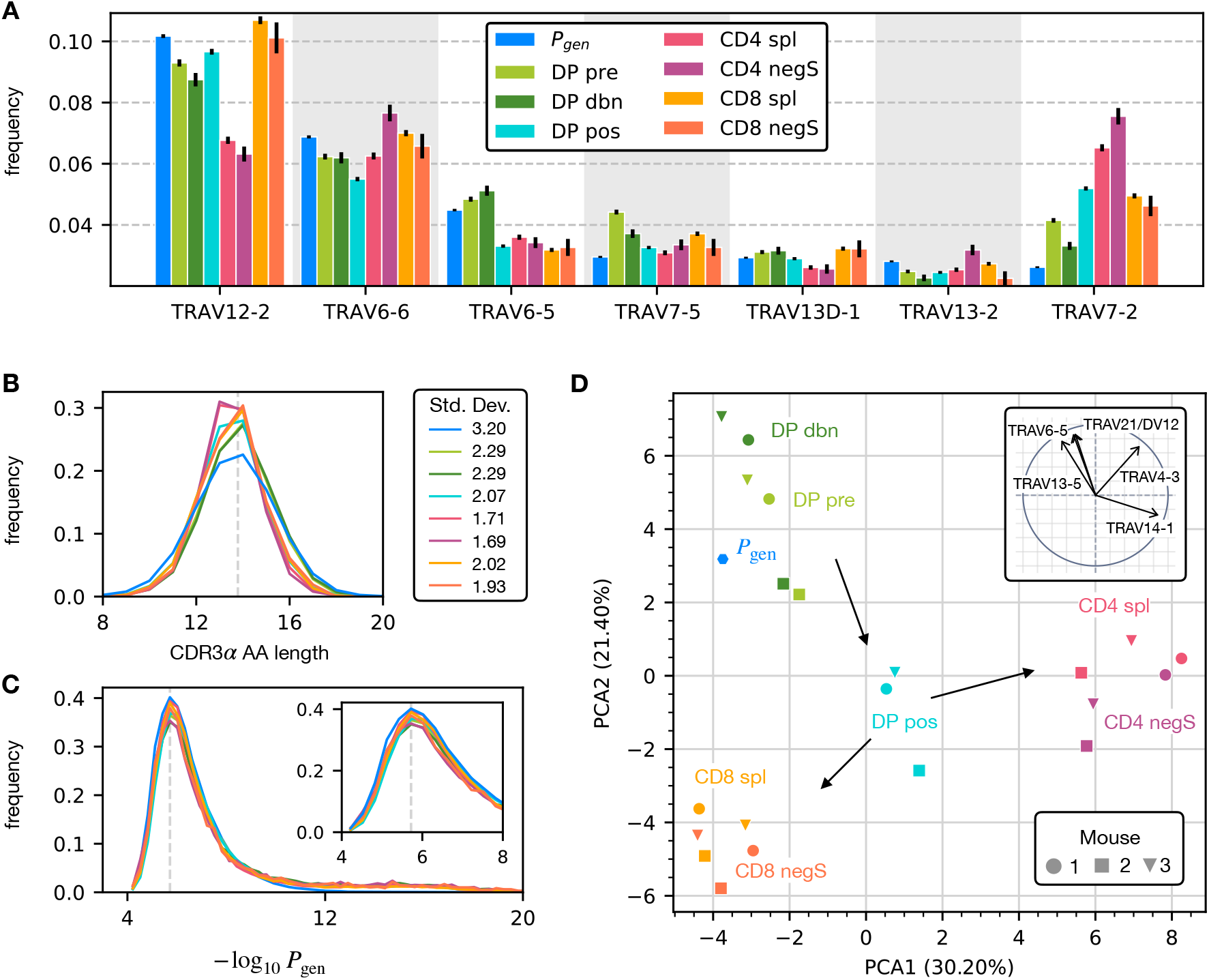
*α* chain sequence properties. The color code is common to all subplots. **A** TRAV gene distribution at different maturation stages compared to the pre-selection model distribution *P*_gen_ (see Fig. S3 for TRAJ). Only the most frequent according to the *P*_gen_ model are reported. **B** CDR3 length distribution of *α* chains. The dashed line is the average CDR3 length from the *P*_gen_ model. Standard deviations are shown at right. **C** *P*_gen_ distribution of *α* sequences at each maturation stage. Insert: blow up of the peak of the distribution. **D** Principal component analysis of the TRAV gene distribution at each maturation stage. Insert: projection on the principal axis of the five most abundant TRAV genes (see Materials and Methods).

For both chains, CDR3 amino acid length of SP CD4+ and CD8+ has a sharper distribution compared to earlier maturation stages (DP), see Figs. 2B and S6B). This has previously been interpreted as a signature of selection due to structural constraints on the pMHC-TCR complex [20–22]. The overall distribution of TCR generation probabilities, *P*_gen_, does not change from the pre-selection and post-selection thymic stages to the mature peripheral SP repertories (Fig. 2C, S6C), consistent with previous reports comparing thymic and peripheral repertoires [23].

The repertoires from different maturation stages cannot be distinguished by any one individual feature discussed above. However, Principal Component Analysis (PCA) on the TRAV gene usage distributions in individual mice at different stages identified clusters of related cell types (Fig. 2D). The DP Nur77- populations cluster with the pre-selection model, the SP CD4+ and CD8+ populations form distinct clusters, and the DP pos Nur77+ cells, which we hypothesise are cells in the process of positive selection, occupy an intermediate position between these three clusters. This pattern is consistent with the known developmental trajectory as illustrated by the arrows in Fig. 2D. PCA of TRAJ usage also shows similar clustering patterns (Fig. S4). The PCA of TRBV and TRBJ usage also discriminates between SP CD4+ and CD8+ populations, and from the pre-selection populations, although the overall pattern is less clear (Figs. S6D and E).

In summary, the effects of selection impose subtle changes on the pattern of TCR variable gene usage, which cannot be adequately captured by looking at any single V or J gene, but only by a combination of features.

V and J gene usage, and CDR3 length are coarse grained measures of a TCR repertoire. We therefore explored whether the repertoires of different maturation stages could be linked to more precise features of the TCR sequence, in particular incorporating the sequence of the CDR3. We encoded each TCR as a sparse {0, 1} binary vector 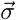 which captures V gene, J gene and CDR amino acid sequence (for details see Materials and Methods). We then trained a logistic regression model on the set of 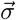 from repertoires of different subsets. We trained and tested the classifier to distinguish pairs of repertoires from different subsets. The classifier achieved only moderate Area Under the Curve (AUC) of the Receiver Operating Characteristic (ROC) scores (Figs. 3A for TRA, and S17 for TRB), in agreement with previous studies [24, 25]. In particular the classifier was not able to reproducibly distinguish between putative selected and non-selected repertoires at the single TCR level. However, the classifier achieved much better performance on *sets* of TCRs taken from different populations. For example, combining the prediction from 30 TCR sequences, the classifier achieved an AUC score of > 0.85 when distinguishing CD4 spl and CD4 negS repertoires (Fig. 3B). Thus statistical properties of a repertoire can distinguish it from another repertoire, even when the feature distributions of individual TCRs are largely overlapping.

**FIG. 3:**
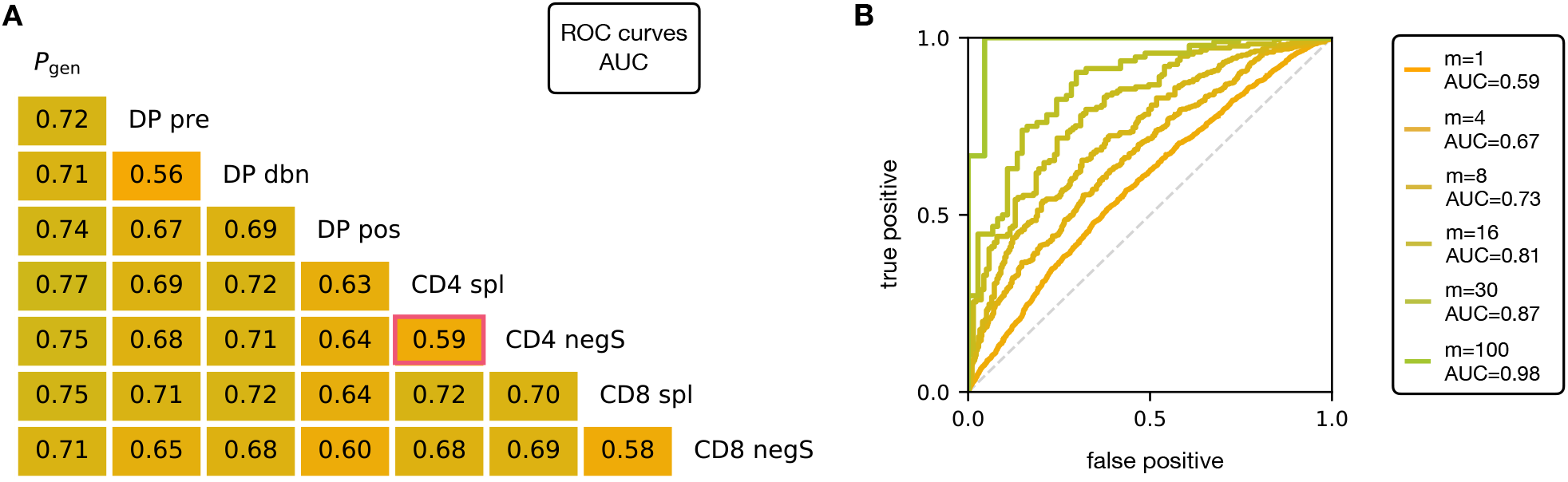
**A** Area under the curves (AUC) values computed from Receiver Operating Characteristic (ROC) curves of linear classifiers of TRA between two subsets. The training/testing set is a random subsample containing 70%/30% of the full dataset at a given maturation stage. Most values are moderately above chance (AUC= 0.5), implying that single cells cannot be classified from their TRA alone. **B** ROC curves for classifying a group of *m* sequences from the same maturation stage, between CD4 spl and CD4 negS (red frame in Fig. 3A), showing that collective discrimination is possible. See Fig. S17 for analogous analysis on TRB.

### Selection models and *n*-grams capture the relations between the stages of thymic development

A number of studies have highlighted the importance of short amino acid motifs (*k*-mers or *n*-grams) within the CDR3 sequence in determining TCR specificity [26–28] (see Fig. 4A). Specifically, *n*-grams can be used to reduce the dimensionality of the TCR space, while capturing amino acid correlations or patterns which might play a role in anitgen recognition. We therefore counted the frequency of *n*-grams in each repertoire. We excluded from the analysis the most conserved regions (the first two positions in the CDR3 that are usually a cysteine and alanine, and the last one, typically a phenylalanine). We then used these *n*-gram frequency distributions to calculate the diversity of the repertoire as quantified by the Shannon entropy *S* (see Materials and Methods). In practice, the Shannon entropy is computationally too expensive to calculate exactly for very large data sets, and we therefore restricted our analysis to *n*-grams of length 4 or less, using the approximate Nemenman-Shafee-Bialek (NSB) entropy estimator [29] to correct for finite sampling bias (see Materials and Methods). This estimator was checked for convergence as a function of sample size (Fig. S7), and shown to outcompete alternative entropy estimators on synthetic data (Fig. S8).

**FIG. 4:**
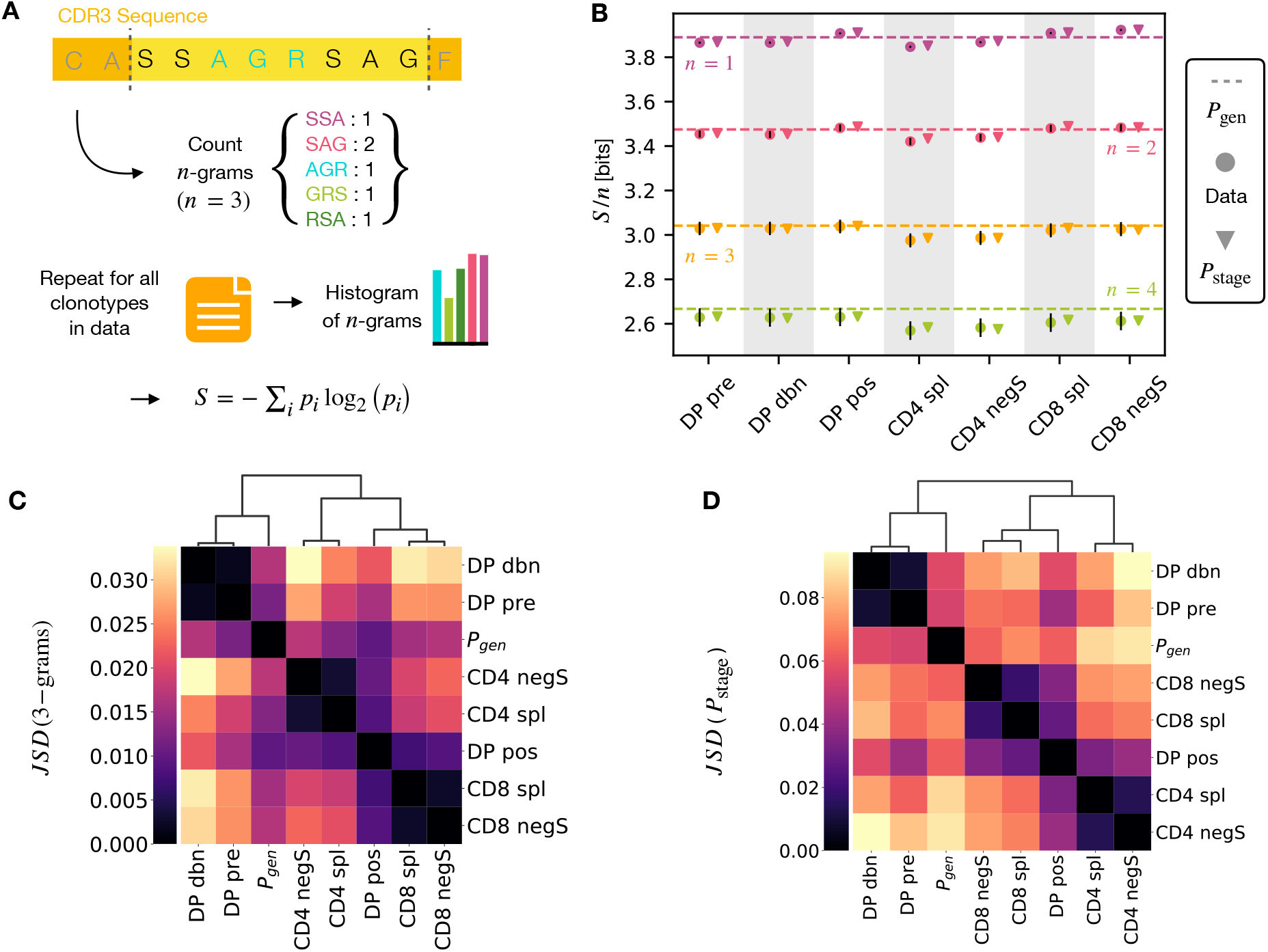
A *n*-gram definition. We count how many times *n*-gram amino acid subsequences are seen in the CDR3 across a repertoire. **B** Shannon entropy *S* of the *n*-gram distributions normalized by *n* for the maturation stages. The entropy is estimated with the Nemenmann-Shafee-Bialek [29] estimator and it is expressed in bits. **C** Jensen-Shannon divergence between the 3-gram distributions computed from the selection model *P*_stage_ on synthetic repertoires. Dendrogram are computed with the Ward method (see Materials and Methods). **D** Jensen-Shannon divergence for the full *P*_stage_ selection model using *P*_stage_.

Once we had generated a set of entropy measurements based on *n*-gram frequencies for each different repertoire, we compared the data-derived entropy measurements with the prediction of a simple generative model of each repertoire which treated each feature of each TCR (V gene, J gene and each CDR3 amino acid) as independent. Taking the set of TCR vectors 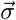 we fitted a set of parameters *E*_stage_ by maximising the posterior probability over all of the TCRs for each repertoire separately 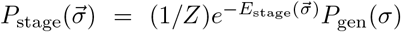, where *P*_gen_(*σ*) are the pre-selection generative probabilities for all the TCRs, 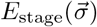 is a linear function of the features [30, 31], and *Z* is a normalization factor (Fig. 1E and Materials and Methods). The enrichment factors 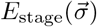 encode the intuition that due to selection, a given TCR in a given repertoire is seen with higher or lower frequency than expected by the pre-selection generation model. Once we had learnt the enrichment factors for each repertoire, we used the resulting model to generate *in silico* synthetic repertoires of 3 × 10^6^ TCRs, and recalculated *n*-gram frequency distributions and entropy estimates for each synthetic repertoire.

The comparison of the estimated entropy for each *n*-gram length, and each subpopulation of T cells, using both directly data-derived and model-derived repertoires is illustrated for TRA (Fig. 4B) and TRB (Fig. S9) chains. The maximum possible entropy if all 20 amino acids were used uniformly would be *S/n* = log_2_ 20 ~ 4.3 bits. Both the observed and model-derived entropies are less than this maximum even for single amino acids (*n*-grams of length *n* =1), and decrease further with *n*-gram length (see Fig S10). This reflects strong selection on the abundance of individual amino acids, and strong correlations between amino acids within the CDR3 which are observed in all CDR3 repertoires, and are captured by the frequency distribution of the longer *n*-grams. Two additional important points can be observed. First, the entropy of the repertoires after selection and lineage commitment (in the single positive populations) is less than the earlier pre-selection DP repertoires, which match closely the entropy of the pre-selection generative model (shown by the dotted line for each *n*-gram length). This decrease becomes more evident with longer *n*-gram length (the circles lie below the dotted lines). Thus, as predicted, selection does impose some decrease in repertoire diversity, although this is a much smaller effect than the decrease in diversity imposed by the generation process itself. The second key observation is that the entropy calculated directly from *n*-gram frequency in the data is very similar to that of the synthetic repertoires generated using the linear generative models in which individual TCR amino acids are treated as independent variables. Thus, at least at the level of diversity, there is no evidence that selection at any step involves complex sequence motifs, or amino acid interactions, which would not be captured by the linear model. Because of sequencing errors, the entropy of *n*-grams is systematically over-estimated in the data. To estimate and correct for this bias, we measured the error rate from the data, provided as a byproduct of the IGoR training procedure [18]. We used this rate to produce synthetic sequences with simulated sequencing errors. The difference in *n*-gram entropy between error-prone and error-free sequences was then applied as a subtractive correction factor to the data. The result shows excellent agreement between the data and the prediction from the selection model (Fig. 4B), validating the hypothesis of linear selection factors.

We looked in more detail at the *n*-gram (*n* = 3) distributions derived by the linear selection models for the different maturation stages. A plot of the Jensen-Shannon divergences (JSD) between all pairwise comparisons largely recapitulated the expected relationships between the subsets, with DP pre and DP dbn clustering with the pre-selection generative model, while the single positive CD4 and CD8 populations clustered separately, and DP pos have an intermediate position (Fig. 4C, S11 for TRA, and Fig. S12 for TRB).

We can go beyond *n*-grams and use the subset-specific *P*_stage_ models to predict the entropy of the full sequence (Materials and Methods), shown in Fig. S13 for TRA and Fig. S14A for TRB. This entropy is substantially reduced from generation to the DP stages, and further reduced in the single positive stages, especially in CD4+ subsets. We also computed the JSD of the distributions *P*_stage_ between subtypes (Fig. 4C for TRA and S14B for TRB). These JSD showed similar patterns as with *n*-grams, except for CD8+ spleen cells showing more similarity to the *P*_gen_ distribution in TRB. Note that the absolute values of the entropies and JSD are larger, since they include information about longer sequences, with additional V and J gene usage information.

In summary, we fitted the data with a set of stage-specific generative models based on linear weighted combinations of TCR sequence features. The repertoires generated by this model accurately estimate the sequence and *n*-gram entropy derived directly from the data, and generate repertoires which differ in a small but reproducible manner from each other. The magnitude of these differences reflect the expected developmental relationships between the different populations.

### Discriminatory power of thymic selection

The stage-specific enrichment factors in the generative models described above can be considered as capturing the combination of features which drives a particular selection step. A prediction of this idea is that, at each selection point, the TCRs which are selected and those which are not would have a distribution of model probabilities (*P*_stage_) which are anti-correlated. For example, a TCR that is present in the DP pos repertoire but “forbidden” from the CD4 repertoire (e.g. because of cross-reactivity to a Class II self pMHC) would be expected to have a large positive *P*_DP pos_ and a *P*_CD4 spl_ ≈ 0, reflecting the large enrichment factor between these two populations. A toy example illustrating this idea is illustrated in (Fig. 5A, left). We consider a simple model in which TCRs are selected according to their CDR3 length into a “long” population with probability 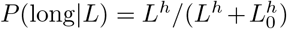, with *L*_0_ = 13 and *h* = 5, and into a “short” population otherwise. We apply this selection process *in silico* to *P*_gen_-generated TCRs, and fit a separate *P*_population_ model on the synthetic sequences found in each subset. We then calculate 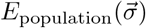 for each TCR in each subset according to both models, and plot these values against each other. The distribution of enrichment strengths according to the two models are clearly anti-correlated. In other words, if a TCR is more likely to be classified as a “long” sequence, it is in general less likely to be classified as a “short” one. Interestingly, however, the enrichment strengths distributions from the two models are significantly overlapping. As a result, attempts to classify individual TCRs according to their enrichment strengths is poor, AUC ~ 0.7 (Fig. 5A, bottom).

**FIG. 5:**
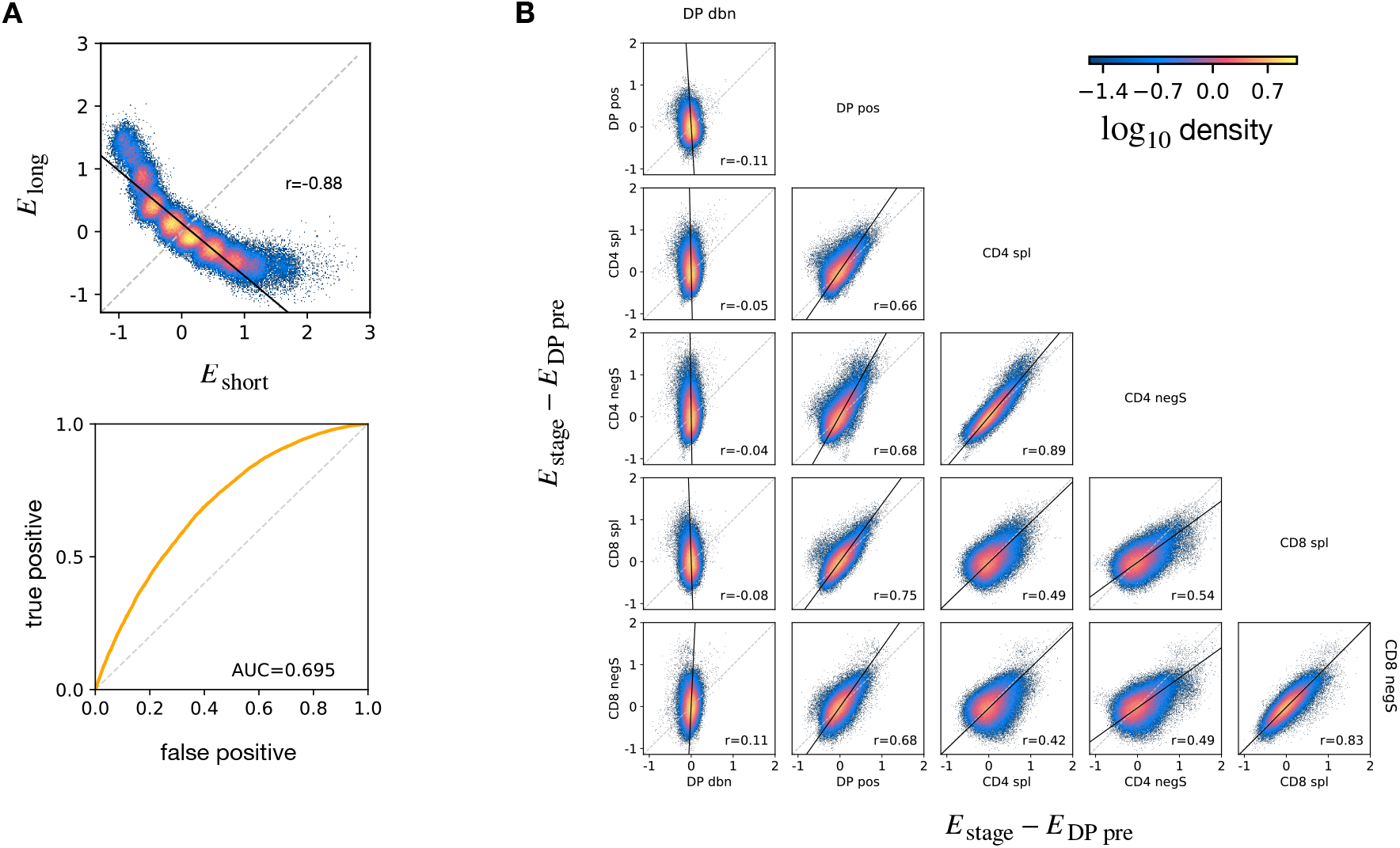
**A** Synthetic example of soft discrimination between “short” and “long” CDR3, where sequences are randomly assigned into either of the two populations with a bias that depends on their CDR3 length. The density scatter plot shows a clear anti-correlation between the selection energies learnt from these two populations. Yet, sequence classification is error-prone, as quantified by the relatively low AUC of the ROC. **B** Density scatter-plots of TRA sequences comparing the selection energies learnt at two different stages. The DP pre energy, which encodes background selection common to all stages, is subtracted. The black line is the direction of the major eigenvector of the points moment matrix. The value *r* reported in each plot is Pearson’s correlation coefficient (see Materials and Methods). See Fig. S15 for analogous analysis on TRB.

We extended this approach to look for relationships between enrichment strengths for TCRs at different developmental stages (Fig. 5B). Since all cells pass through a DP pre-selection stage, we can consider *P*_DP pre_ as a common background distribution for all the other thymic stages. We therefore considered differential enrichment parameters *E*_stage_ – *E*_DP pre_, a linear operator which predicts whether a sequence is more or less likely to be present in a particular developmental stage as compared to DP pre. We generated a set of sequences using the pre-selection *P*_gen_ model, and then computed enrichment factors for each TCR according to all the stage specific models, relative to the DP pre. The set of pairwise correlations between enrichment values for the different populations are shown in (Fig. 5B for TRA and Fig. S15 for TRB). In contrast to the toy model illustrated above, most plots showed a positive correlation between enrichment values for two models. Thus a common dominant selection process is driving the repertoire shift between the DP pos and all subsequent stages, which dominates the impact of individual stage specific selection processes. The exception was the DP dbn repertoire, which showed a narrow distribution of values, which was uncorrelated to any other subset. This is consistent with the DP dbn repertoire containing a random sample of the DP pre repertoire, unrelated to its TCR sequence. Surprisingly the spleen SP and the thymic negS populations were highly correlated for both CD4+ and CD8+ cells (*r* = 0.89 for CD4 spl vs CD4 negS, *r* = 0.83 for CD8 spl vs CD8 negS). There was therefore no evidence of selection pressure operating on TCR sequence to distinguish these two populations. The correlation between the CD4+ and CD8+ subsets was weaker although still positive (*r* ~ 0.42 - 0.54), suggesting that the selection pressures operating on the two populations are distinct, although not mutually exclusive. Similarly, inspecting single amino acid usage in terms of the model marginals (see Eq. (3)) we do not observe any striking signal (Fig. S16).

In summary, the TCR enrichment value distributions differ between different thymic populations, but do not show evidence of dominant exclusive sequence-based selection operating at any step of the selection process.

## III. DISCUSSION

Thymic selection is often portrayed as a simple discrimination process that eliminates TCRs capable of strongly binding any self-peptide, while promoting TCRs that bind them weakly. This simple picture is very appealing, but is difficult to reconcile with the known biology of T cell development. Murine thymocytes spend maximally 4-5 days in the thymus [32]. Within this brief window, each T-cell is estimated to encounter 500 to few thousand self-peptides [7, 33], while the number of presentable self peptide-MHC complexes is estimated from the peptidome to be ~ 5 · 10^5^ – 5 · 10^6^ [33]. If each T cell only meets a small fraction of the self-peptidome, many self-reactive T cells will escape negative selection [34] and the mature repertoire will contain many self-reactive cells. A further implication is that no sequence feature will unambiguously distinguish TCRs from pre and post-selected repertoires. Many efforts have been made to connect TCR sequences to peptide recognition [25, 35, 36]. However, these approaches cannot yet be used to define the target peptidome of entire repertoires. Here we take the complementary approach, by looking for TCR sequence features that are linked to positive or negative selection.

Although there has been a lot of work on understanding and modeling thymic development [2, 4] our study presents the first comprehensive analysis of TCR repertoire of different developmental stages of thymic maturation. In addition, by incorporating a reporter for the activation marker Nur77, which is activated during thymic selection, and an early marker of apoptosis, Annexin V, we were able to identify subpopulations during the process of positive or negative selection. We were therefore able to directly compare the repertoires at the major stages of thymic selection.

We examined the repertoires from two perspectives. In the first part of the paper, we compare statistical properties of the sequences of the repertoires using features of different dimensionalities, which include V gene, J gene and CDR3 length frequency distributions, and individual CDR3 sequences represented as sparse {0, 1} binary vectors. The analysis incorporated both coarse-grained (V, J and CDR3 length) and fine-grained (individual CDR3 sequence) features, and the results were remarkably consistent. No single feature adequately discriminated between any pair of repertoires. Combination of features when averaged across a repertoire did show subtle but reproducible differences between repertoires, which could be used to discriminate between subpopulations using both unsupervised (PCA) and supervised (logistic regression) analysis. Furthermore, the difference between these statistical parameters captured the known developmental trajectory of thymic development, illustrated schematically in Fig. 1D. Interestingly, the smallest distances observed were between mature CD4 or CD8 cells, and their thymic SP negatively selected (negS) counterparts. Thus, either negative selection of single positives is only weakly associated with TCR sequence properties, or the Annexin V staining does not adequately capture the negatively selected population.

Although the statistical properties of the repertoires differed between subpopulations, it was not possible to classify individual TCRs at high accuracy. Instead, learning the collective properties of at least a few dozen TCRs was required in order to achieve good discrimination between repertoires. These observations are compatible with previous models emphasising the importance of collective, rather than individual T cell behaviour. Butler et al [33], proposed that a minimum number *t* of T-cells must collectively recognize a peptide to trigger a response, proposing quorum sensing as a mechanistic explanation of this collective decision making. Recent experiments have confirmed that quorum sensing between TCRs can occur, mediated via cytokine signaling [37], and estimating a minimum quorum size of activated T cells to be ≳ 30. This number is in agreement with our observations (Fig. 5B) that ~ 30 TCRs increases the AUC of correctly classifying these cells as CD4+ spleen vs CD4 negS from ~ 0.59 to ~ 0.87. In the context of self non-self discrimination, our results suggest that thymic selection imposes only a rather weak selective pressure on the repertoire. Even a subtle depletion, rather than complete elimination of non-self TCRs, however, may translate into robust self/non-self discrimination at the population level.

In the second part of the study we explore in more detail whether we can discover any evidence that thymic selection depends on specific sequence motifs (i.e. a strong correlative structure between CDR3 amino acids). For this purpose, we build on our previous work which have established a framework for the development of generative statistical models of repertoire generation, based firmly on a mechanistic understanding of TCR generation and selection. Specifically, we construct models which incorporate only linear combinations of CDR3 sequences to capture the selective process which can transform one repertoire into another. These models produce an “enrichment factor” for each TCR which estimates its relative likelihood of being in a particular stage-specific population. Intuitively, one can consider these factors as capturing the probable enrichment or depletion of a TCR with a particular sequence when comparing two repertoires. We demonstrate that these linear models effectively capture the progressive decrease in repertoire diversity which we observe in the preselected DP to the SP transition. They also effectively capture the known developmental relationships between the thymic subpopulations. Thus we find no evidence that complex non-linear amino acid sequence interactions are required to explain the observed changes in repertoire observed in our data. We also compared the distributions of enrichment factors between populations. We demonstrate that, contrary to the predictions of a strong binary selection model, we do not observe any negative correlation between enrichment factor distributions between selected and non-selected repertoires. Instead, we observe a set of positive correlations, revealing a dominant conserved selection process spanning the developmental stages between pre-selection DP and mature SP. Consistent with the clustering data discussed above, we find strong correlation between the enrichment factor distributions of mature SP and thymic negatively selected population, and no evidence of binary selection between these two populations.

In conclusion, we report a comprehensive analysis of the TCR repertoire at various stages of thymic development. We then combine data-driven and model-based analysis of these repertoires. Our conclusions are incompatible with a model of thymic developments which involves a sequence of clear-cut binary selection processes, based on TCR sequence features. Rather, our data suggest a probabilistic fuzzy decision making process at each selection step. We propose that this model is compatible with robust self/non-self discrimination, if T cell responses to antigen are governed by collective quorum based decision making. Further experimental and theoretical work is required to test these hypotheses, which have fundamental implications for strategies to modulate the immune response for prophylaxis or therapy of human disease.

## IV. MATERIALS AND METHODS

### Animals

The experiment was carried out using three 6-weeks old male inbred Nur77-GFP/Foxp3-mCherry (C57BL/6 background) [38]. The cross was bred and maintained at the Weizmann institute. All animals were handled according to Weizmann Institute’s Animal Care guidelines, in compliance with national and international regulations.

### Sample preparation and T cell isolation

Thymocytes and splenocytes were isolated from Nur77-GFP/Foxp3-mCherry 6-weeks old mice. Erythrocytes were removed by hypotonic lysis in ammonium chloride. Thymocytes were stained with fluorescent antibodies, and sorted using a flow cytometer as described below. Splenic CD4 and CD8 cells were purified in two steps: (1) CD4+ positive selection (CD4 (L3T4) MicroBeads, mouse, # 130-117-043, Miltenyi) to generate the “CD4 spl” samples (2) the negative cells fraction were further selected for the CD8+ positive cells (CD8a (Ly-2) MicroBeads, mouse, # 130-117-044, Miltenyi Biotec) to generate “CD8 spl” samples.

### Flow cytometry analysis and cells sorting

The following fluorochrome-labeled mouse antibodies were used according to the manufacturers’ protocols: PerCP/Cy5.5 anti-CD4, PB anti-CD8, PE/cy7 anti-CD3, APC annexinV (Biolegend). UV LIVE/DEAD™ (ThermoFisher Scientific, # L23105). Labelled cells were sorted on a SORP-FACS-AriaII using a 70 μm nozzle to 5 populations (see Table 1). Cell counts are reported in Table S2. Cells were analyzed using *FlowJo* (Tree Star) software.

**TABLE 1:** Cell sorting based on fluorochrome-labeled mouse antibodies.

| Sample\Marker | CD4 | CD8 | CD3 | AnnexinV | Nur77 |
| --- | --- | --- | --- | --- | --- |
| DP pre | + | + | + | - | - |
| DP pos | + | + | + | - | + |
| DP dbn | + | + | + | + | - |
| CD4 negS | + | - | + | + |  |
| CD8 negS | - | + | + | + |  |

### Library preparation for TCR-seq

All libraries in this work were prepared according to the published method [39], with minor adaptations as described below. Briefly, total RNA was extracted from each of the seven populations using RNeasy Micro Kit (# 74004, Qiagen) and cleaned from excess DNA with DNAse 1 enzyme (# M6101, Promega). RNA samples were reverse transcribed to cDNA (SuperScript™ III, # 12574026, Invitrogen) using primers for the mouse alpha chain (mAlpha_RC2) and for the mouse beta chain (mBeta_RC2) (see Table S1). Following reverse transcription the samples were purified on minielute spin columns (# 28004, QIAGEN). The cDNA was ligated to an oligonucleotide containing a unique 12 basepair molecular identifier (UMI) (6N_I8.1_6N_I8.1_SP2, see Table S1) using T4 RNA ligase (M0204S, NEB). Ligation products were purified using Agencourt AMPure XP beads (# A63881, BeckmanCoulter). Next, three rounds of extension PCR were executed (using KAPA HiFi DNA Polymerase, KAPA Biosystems) to add illumina sequencing adaptors and Illumina sample indices for multiplex sequencing (see Table S2). The thermal cycler parameters are an initial denaturation step (3 minutes at 95°C) followed by cycles of denaturation (98°C for 20 seconds), annealing (61°C for 15 seconds), and extension 72°C for 30 seconds. The final extension step was at 72°C for five minutes. The lid was maintained at 105° C. After the first round PCR (5 cycles), PCR products were purified using Agencourt AMPure XP beads and split in two, and alpha and beta TCR genes were processed separately in subsequent steps. After the second PCR (8 cycles), PCR products were again purified using Agencourt AMPure XP beads. The final amplification using the adapter sequences P5 and P7 were carried out on a real-time qPCR machine, and the amplification was tracked by the incorporation of SYBR green. The cycler was stopped manually when the fluorescent signal reached a predetermined threshold, thus preventing overamplification.

The final library concentration was measured using Qubit Fluorometric Quantification (ThermoFisher Scientific) and the presence of the correct 600-700 bp product confirmed by electrophoresis on a High Sensitivity D1000 ScreenTape cassette using a 4200 TapeStation System (Agilent). Multiple samples were pooled in equal molarity, and then sequenced using NextSeq 550 (200 bp forward read, 100 bp reverse) (Illumina).

### Pre-Processing and Error Correction for Raw Reads

Data were processed using an in-house pipeline, coded in R. First, UMI sequences were transferred from read2 to read1. Trimmomatic was used to filter out the raw reads containing bases with Q-value ≤ 20 and trim reads containing adaptors sequences [40]. The remaining reads were separated according to their barcodes and reads containing the constant region for α or *β* chain primers sequences were filtered (CAGCAGGTTCTGGGTTCTG-GATG / TGGGTGGAGTCACATTTCTCAGATCCT *α* and *β* chain, respectively), allowing up to three mismatches. To correct for possible sequence errors, we cluster the sequences UMIs’ in two steps; (1) The UMIs with the highest frequency are grouped within a Levenshtein distance of 1 [41]. (2) Out of these sequences, CDR3AA sequences (starting from the most frequent sequence in a group) were clustered using a Hamming distance threshold of 4 [42]. Finally, the UMI of each CDR3 sequence was counted.

### Annotation and Generation Model

From the raw nucleotide reads, we performed a preliminary annotation using the python module *PyIR* (version 1.3.0) [43], which provides a wrapper and parser of the open source software *IgBlast* [44]. We then separated the productive clonotypes from the out of frame reads and/or reads containing stop codons. We define a clonotype as TCRs sharing V genes, J genes, and the same CDR3 nucleotide sequence. If different reads are annotated as the same clonotype in the same dataset, only the read with highest UMI counts is considered.

For our models, we use a reduced set of genes from the IMGT free online repository [45]) in order to have a single allele per gene, preferring functional alleles to open reading frame or pseudo genes. A further reduction is done for the V genes of the α chain, clustering to a single representative all of the those genes that result indistinguishable in the region from the maximum observed V offset for the annotation to the conserved cysteine. Two genes are said to be indistinguishable if the Hamming distance [42] between the considered regions is equal to 0. For each TRAV cluster, we choose as the representative the most frequent gene in the preliminary annotation. In this way we obtain 76 TRAV genes and 51 TRAJ genes for the α chain, 26 TRBV genes, 2 TRBD genes and 14 TRBJ genes for the *β* chain.

In order to infer a generation model we use the open source software *IGoR* [18] on all out-of-frame clonotypes pooled from all maturation stages of all mice. The generation model associates to each *α* (*β*) read a probability *P*_gen_ of being generated through the VJ (VDJ) recombination process. After learning a generation model, we annotate the reads using the most probable alignment scenario using the IGoR software, as the clonotype (V, J gene choice, CDR3 nucleotide sequence) with the highest *P*_gen_ among all possible recombination scenarios.

The PCA was computed in *R* (version 3.6.0) using the function “PCA” from the *FactoMineR* package (version 2.4).

### Statistical Classification

The features are assigned to each α chain as a binary vector 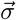, where each entry is equal to 1 if the feature is observed, 0 otherwise. IN this study the set of features is encoded from the “SoniaLeftposRightpos” (from the Python package *Sonia* version 0.0.45) which returns 5033 features: 30 for the CDR3 amino acid lengths, 25 left to right positions for each of the 20 amino acids (500 features), 25 right to left positions (500), the joint V/J gene usage (76 × 51 = 3876) and the independent usage (76 + 51 = 127). Analogously for the *β* we obtain 1434 features (without considering TRBD genes).

To learn the models for the statistical classification of two stages, we first remove all sequences that share the same features between the two sets (i.e. same amino acid CDR3, V and J gene). Then, we balance the size of the sets sub-sampling the larger one so that its size does not exceed 25% of the size of the smaller. Each of the resulting set is divided into a train and a test set by a ratio 70%/30%. The classifiers are learned with linear models, defined by a single layer with binary crossentropy as loss function, binary accuracy as metrics, a sigmoid as activation function and the Adam optimization algorithm for the stochastic gradient descent, coded using the “keras” module from the Python package *tensorflow* (version 2.4.1). We obtained similar performances for the classification task by learning with random forest algorithm as provided by the function “RandomForestClassifier” in the module “ensemble” from the Python package *scikit-learn* (version 0.24.2).

### Selection Model

To learn a *P*_stage_ selection model for each maturation stage, we pooled together the annotated sequences from all mice for the given maturation stage, discarding all clonotypes annotated with non-functional and pseudo genes. We learn a selection model using the open source software *Sonia* for each maturation stage. *Sonia* performs a linear regression over the features of the sequences in the dataset to infer the enrichment ratio between the maturation specific dataset and the generation model. The feature choice for the enrichment model is similar, except for the fact that only independent gene usage is considered, reducing features to 1157 for α chain (1070 for *β* chain). The probability of observing a sequence in a stage is modeled as

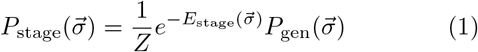

where *Z* is a normalization factor and the energy 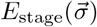 for a sequence showing a set of features 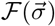 is defined as

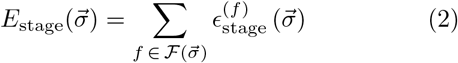

Here *ϵ*^(*f*)^ is a weight associated to the feature *f* and is learnt from data. To look at specific enhanced features between stages *a* and *b* one can obtain the average weights difference from the respective *P*_stage_ models as

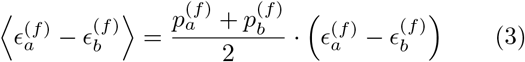

where 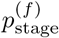 is the marginal associated by the model to the feature.

The limited amount of clonotypes for certain maturation stages precludes using deep neural network based selection models, although we do not expect the conclusions to change with the DNN SoNNia model [25].

### *n*-gram Shannon Entropy Estimation

As a diversity measure we consider the Shannon entropy defined as:

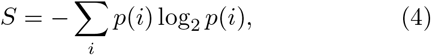

where *p*(*i*) is the probability of finding a clonotype in the data. Since *n*-grams are sampled from 20*^n^* possible motifs, undersampling could bias a naive estimation of the entropy. We overcome this bias by estimating the Shannon entropy using the Nemenman-Shafee-Bialek (NSB) estimator [29]. The NSB estimator is computationally tractable and calculates an estimation error. We implement the entropy and variance estimators as given in [46]. To check for convergence we subsample the clonotypes in the dataset at increasing sizes and estimate the entropy for each sub-sample (Fig. S7). Convergence sets a limit of *n* = 4 due to sample size constraints of the smallest dataset. We repeat the same computation for synthetic repertoires. We verified the NSB estimators better performance for our datasets compared to other non-parametric estimators (Fig. S8), consistently with previous reports [46].

### Full Model Shannon Entropy Estimation

The Shannon entropy in Eq. 4 associated to the full 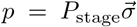 model requires summing over all possible clonotypes 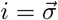. Practically we evaluate the entropy by producing synthetic sequences according to the selection model *P*_stage_ and averaging the value of log_2_ *P*_stage_

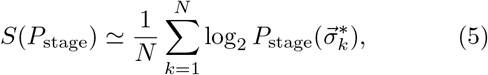

with clonotypes 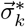 sampled from the *P*_stage_ distribution.

### *n*-gram Jensen-Shannon Divergence

To quantify the distance between two distributions *p_a_* and *p_b_* defined on the same support, we use the symmetric Jensen-Shannon divergence *JSD:*

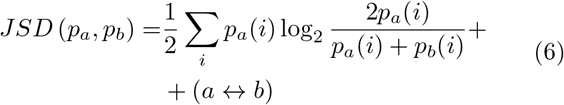

where the sum runs over all possible observables *i* and the term (*a* ↔ *b*) corresponds to the same expression in the first one with a and b inverted. The Jensen-Shannon divergence is bounded between 0 and 1 bits, with JSD = 0 bits if the distributions are identical and a maximal difference of JSD = 1 bit. We use JSD to asses the divergence between *n*-gram distributions and between selection models.

### Full Model Jensen-Shannon Divergence

To compare selection models of complete clonotypes at two maturation stages, the divergence between the *P*_stage_ distribution of model *a* and the model *b* is:

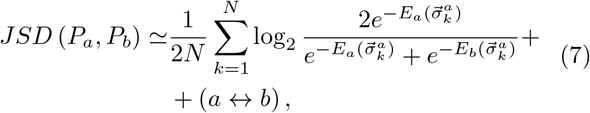

where the clonotypes 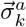 are sampled from the *P_a_* distribution. In Eq. 7, we used the fact that a given sequence has the same background generation probability *P*_gen_ in both selection models.

### Other Software for Statistical Analysis

The Jensen-Shannon dendrograms linkage is computed by the Ward method as provided by the function “linkage”, reordered according to the function “optimal_leaf_ordering”, both from the Python module “cluster.hierarchy” in *scipy* package (version 1.7.3). The Pearson correlation coefficient is computed with the Python function “pearsonr” as contained in the module “stats” in the *scipy* package.

## Code availability

All code for reproducing the figures of this paper can be found at https://github.com/statbiophys/thymic_development_2022.git.

## Acknowledgements

The study was supported by the European Research Council COG 724208 and ANR-19-CE45-0018 ?RESP-REP? from the Agence Nationale de la Recherche and DFG grant CRC 1310 ?Predictability in Evolution?.

---

**FIG. S1:**
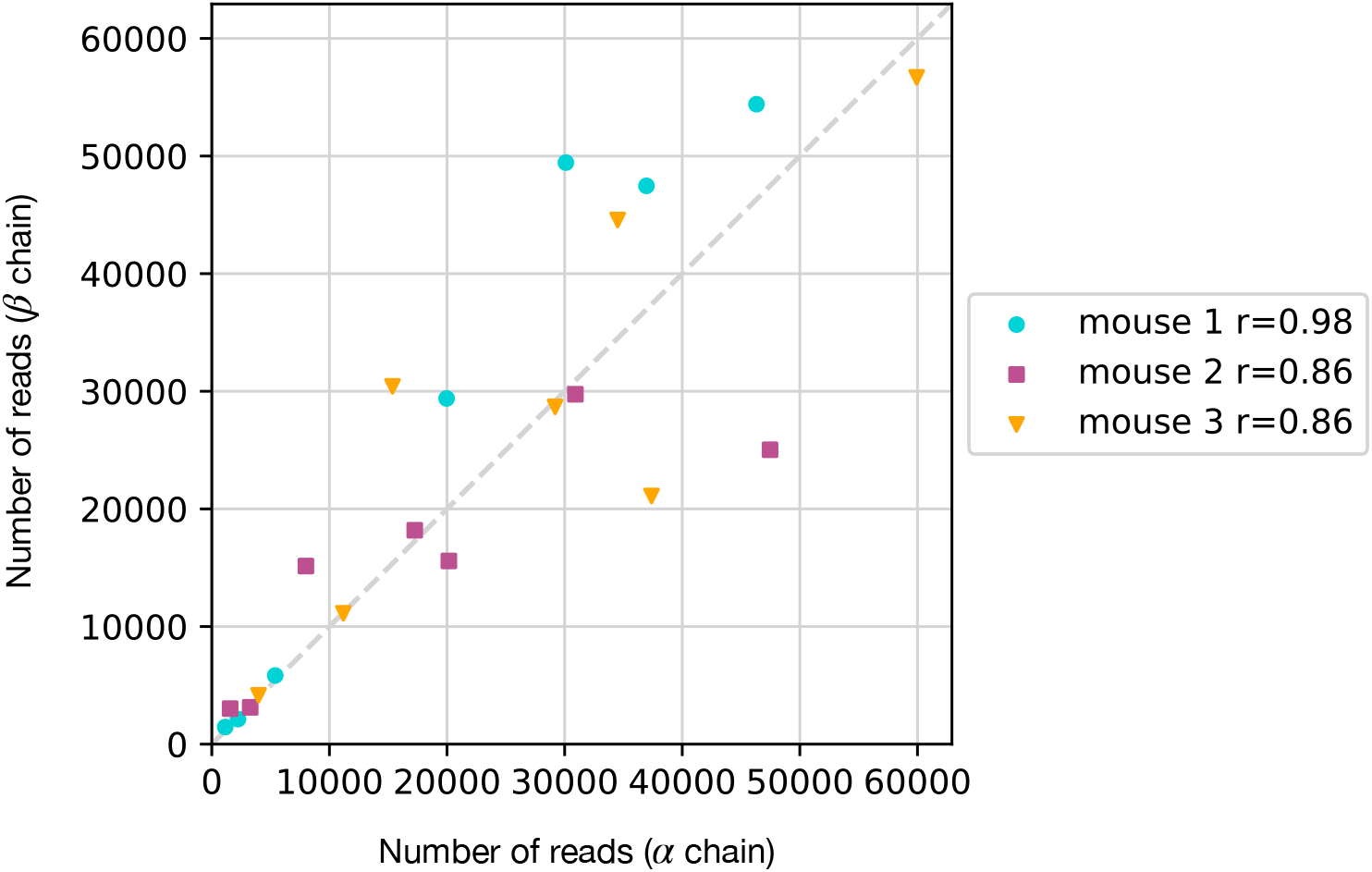
Number of reads for the alpha chain vs the number for the beta chain within the same dataset. In the box is shown the Pearson correlation coefficient.

**FIG. S2:**
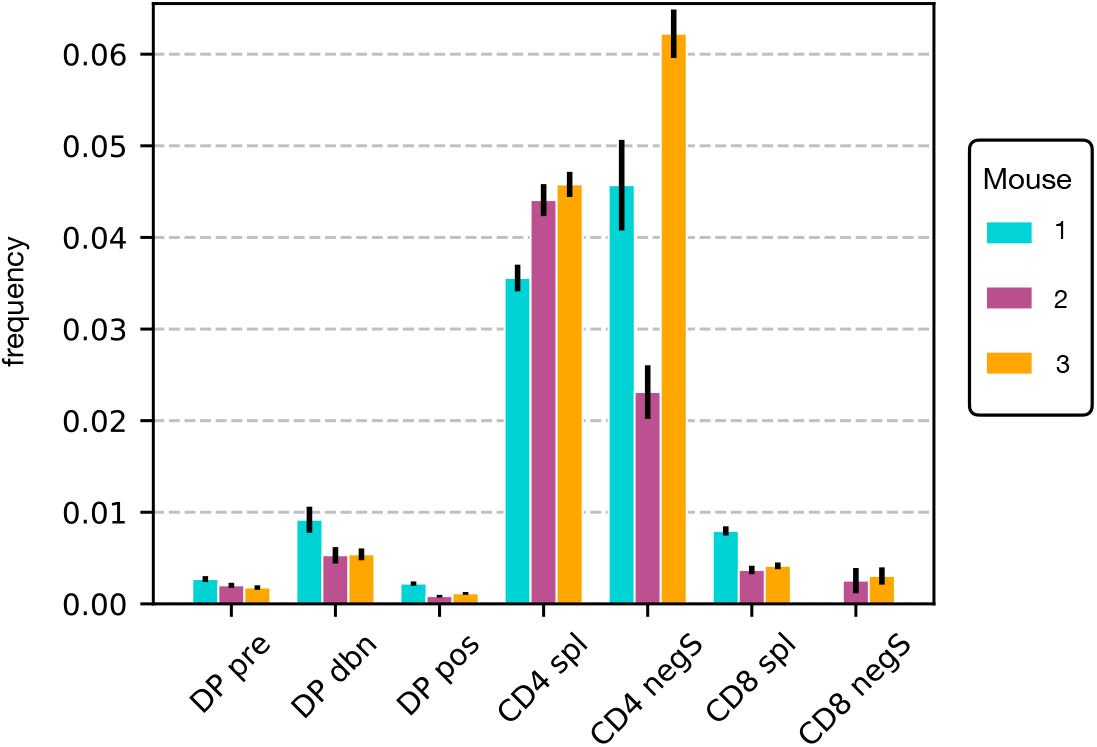
Distribution of iNKT clonotypes for the *α* chain. The relative amount of (TRAV11, TRAJ18) clonotypes is significantly higher for all CD4 stages in all mice. Error bars are computed assuming Poisson statistics for the counts.

**FIG. S3:**
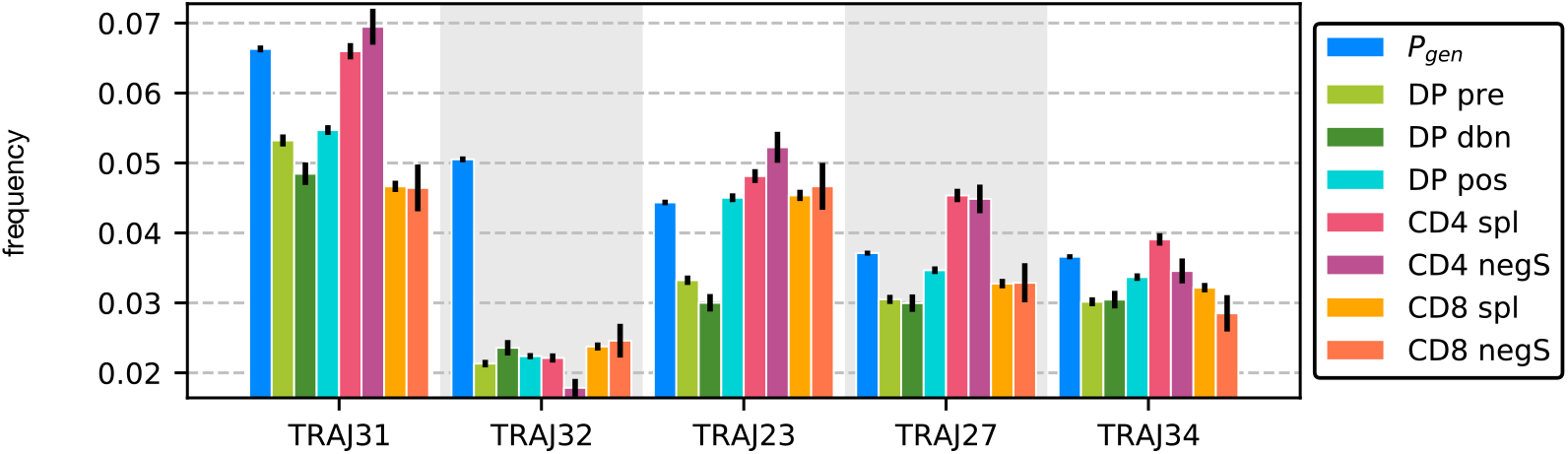
TRAJ gene distribution at different maturation stages compared to the generated distribution.

**FIG. S4:**
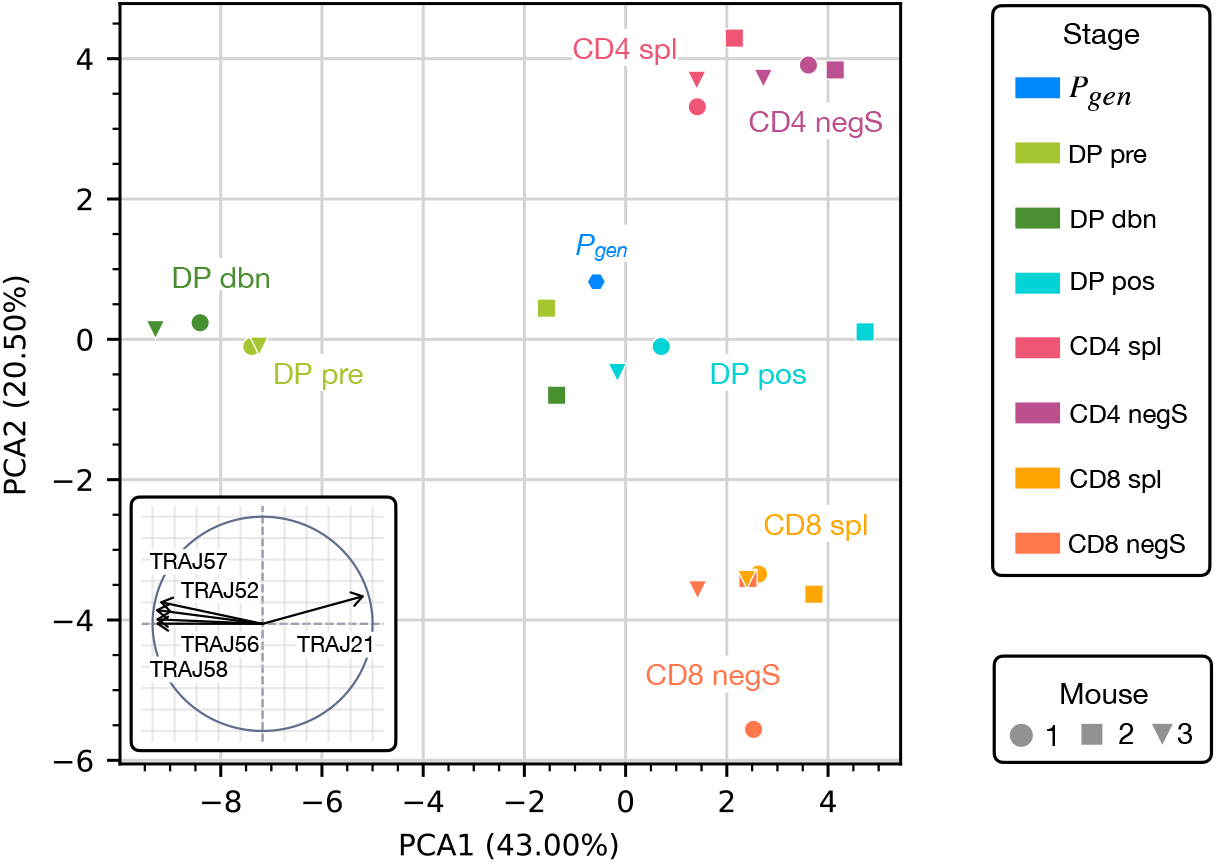
Principal component analysis of *α* chain J gene usage. Similarly as for the TRAV genes (Figs. 2C), DP, CD4 and CD8 maturation stages cluster within the cell types.

**FIG. S5:**
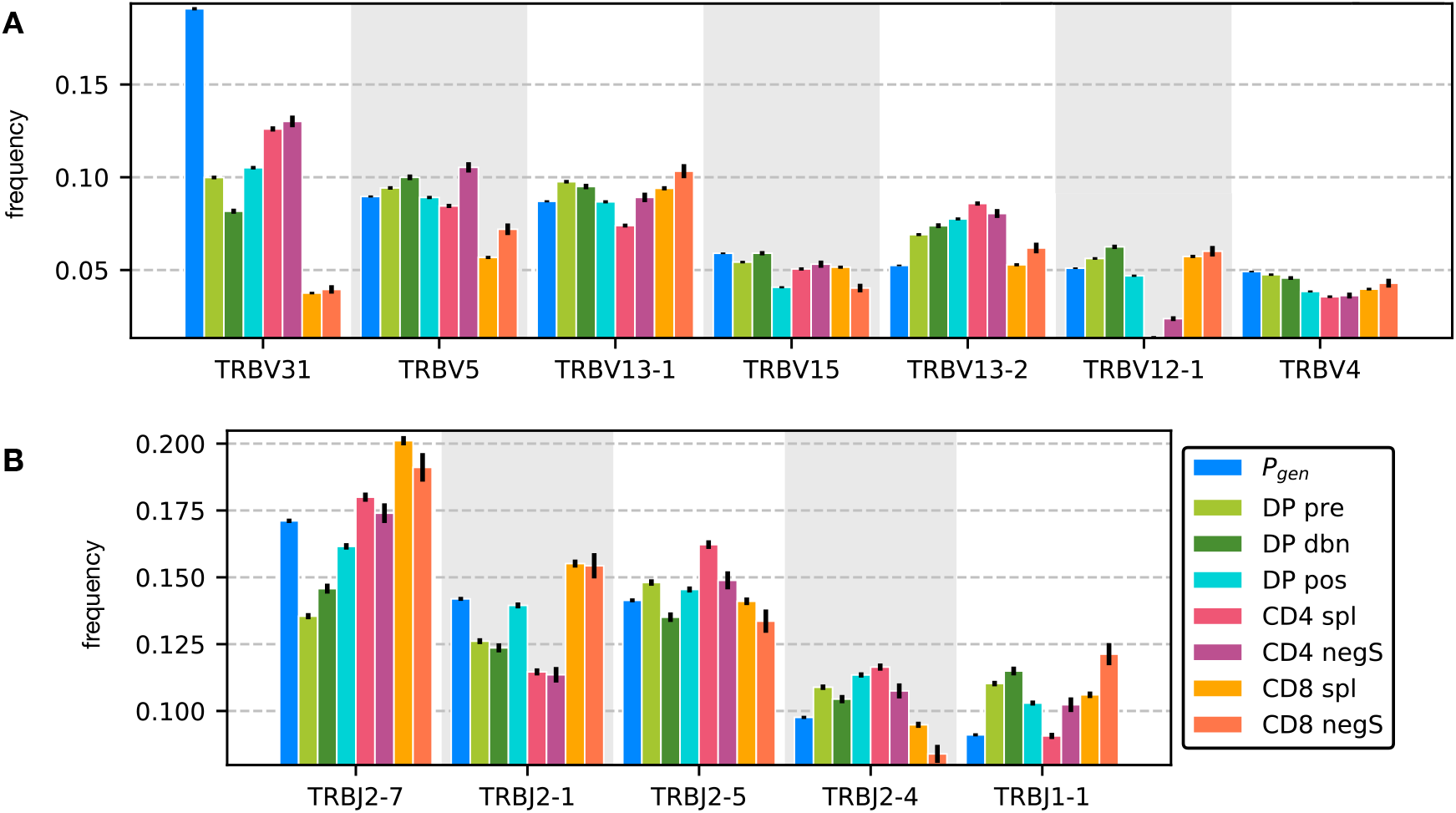
Gene distribution at different maturation stages compared to the generated distribution. **A** The most frequent TRBV genes at different maturation stages. **B** The most frequent TRBJ at different maturation stages.

**FIG. S6:**
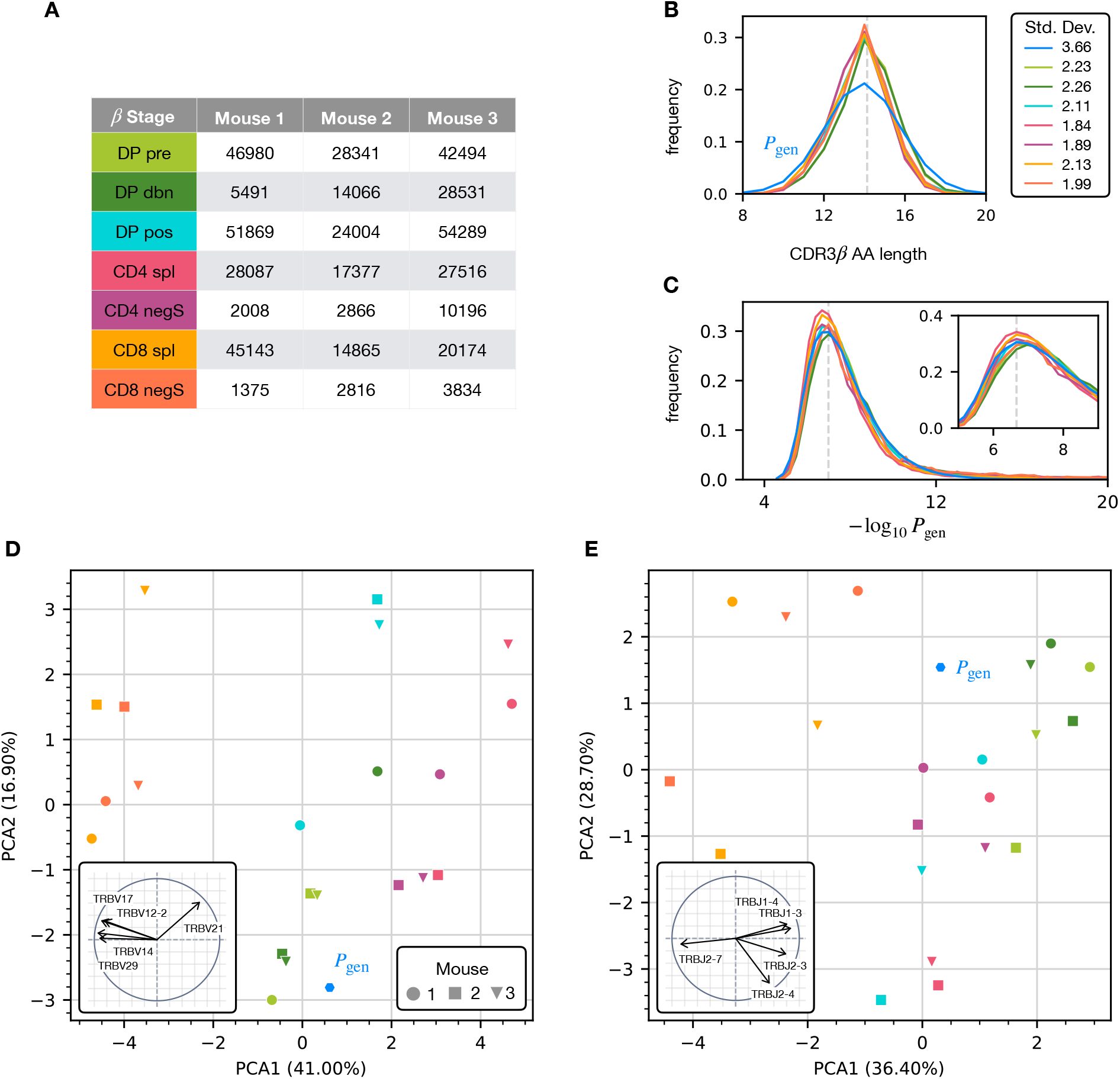
Analysis of the annotated productive *β* clonotypes for the different maturation stages. **A** Numbers of unique productive clonotypes for the *β* chain. The columns provide a colorcode for the figure. **B** Distribution of *β* chain CDR3 amino acid sequence lengths. The CDR3 is defined between the typical cysteine and phenylalanine position. The dashed line represents the average length according to the *P*_gen_ model. **C** Distribution of *P*_gen_ values for *β* clonotypes at each maturation stage. Insert: zoom of the peak (the dashed line) of the distribution of sequences generated with the *P*_gen_ model. **D** Principal component analysis according to the TRBV gene distribution at each maturation stage. Insert: projection on the principal axis of the five most representative TRBV genes. **E** Same as in panel D but for TRBJ.

**FIG. S7:**
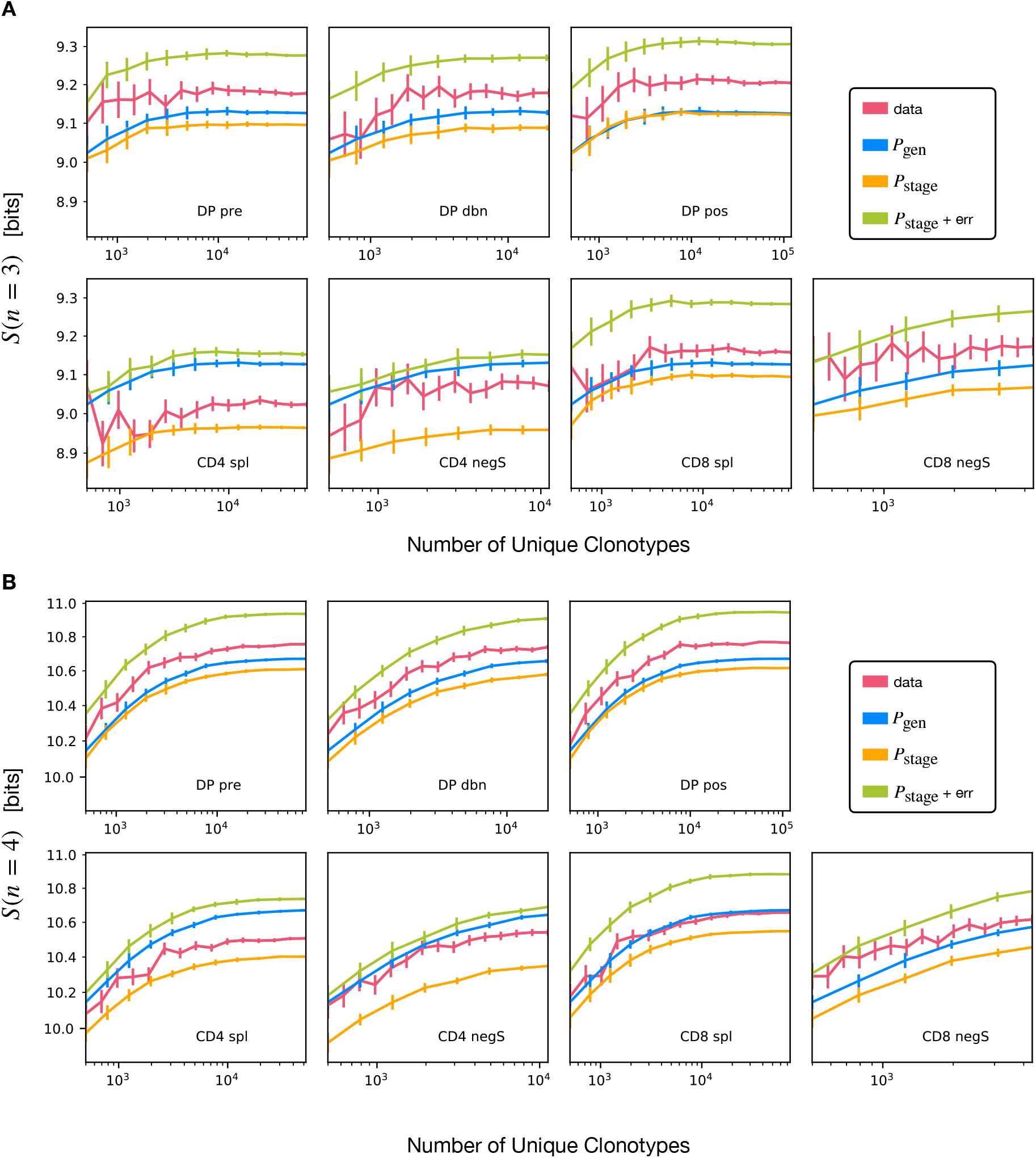
Convergence of the *n*-gram entropy estimation with increasing number of unique clonotypes for *α* chain calculated using the Nemenman-Shafee-Bialek (NSB) estimator. Comparison of estimation from data and from synthetic sequences produced with the generation model *P*_gen_ and the different selection models *P*_stage_, and a *P*_stage_ selection model with nucleotide sequencing error. Error bars for data from the NSB estimator, for synthetic sequences estimated as the standard deviation over different realizations of the simulation. **A** *n* = 3. **B** *n* = 4.

**FIG. S8:**
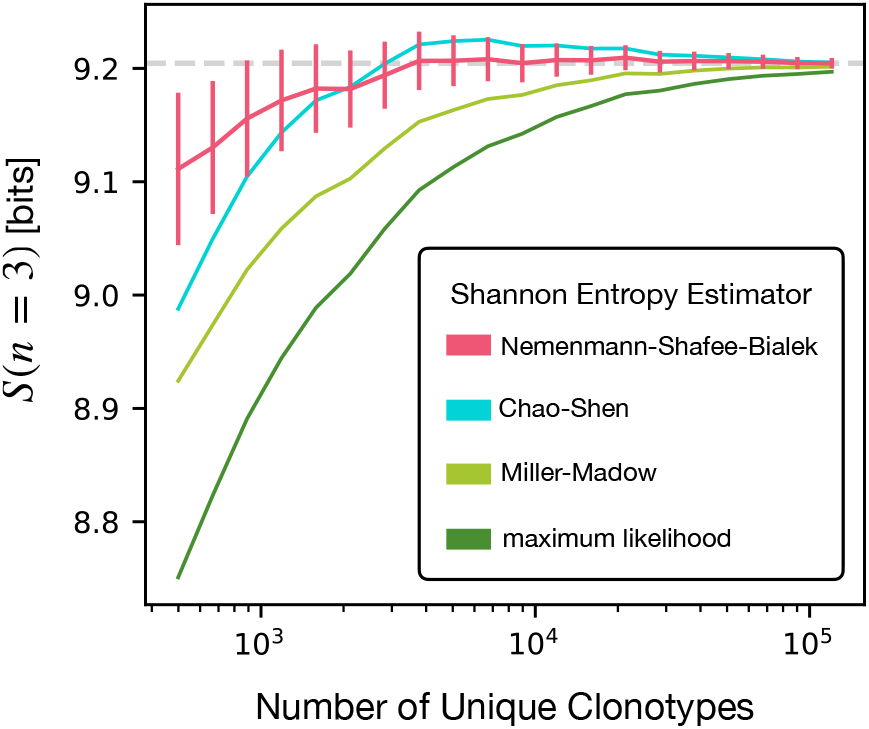
Comparison of different entropy estimators evaluated on subsamples of productive DP pos *α* clonotypes pooled from different mice. Averages over different subsamples of the same size. Estimators: Nemenmann-Shafee-Bialek (see Materials and Methods) [29], Chao-Shen [47], Miller-Madow [48] and maximum likelihood.

**FIG. S9:**
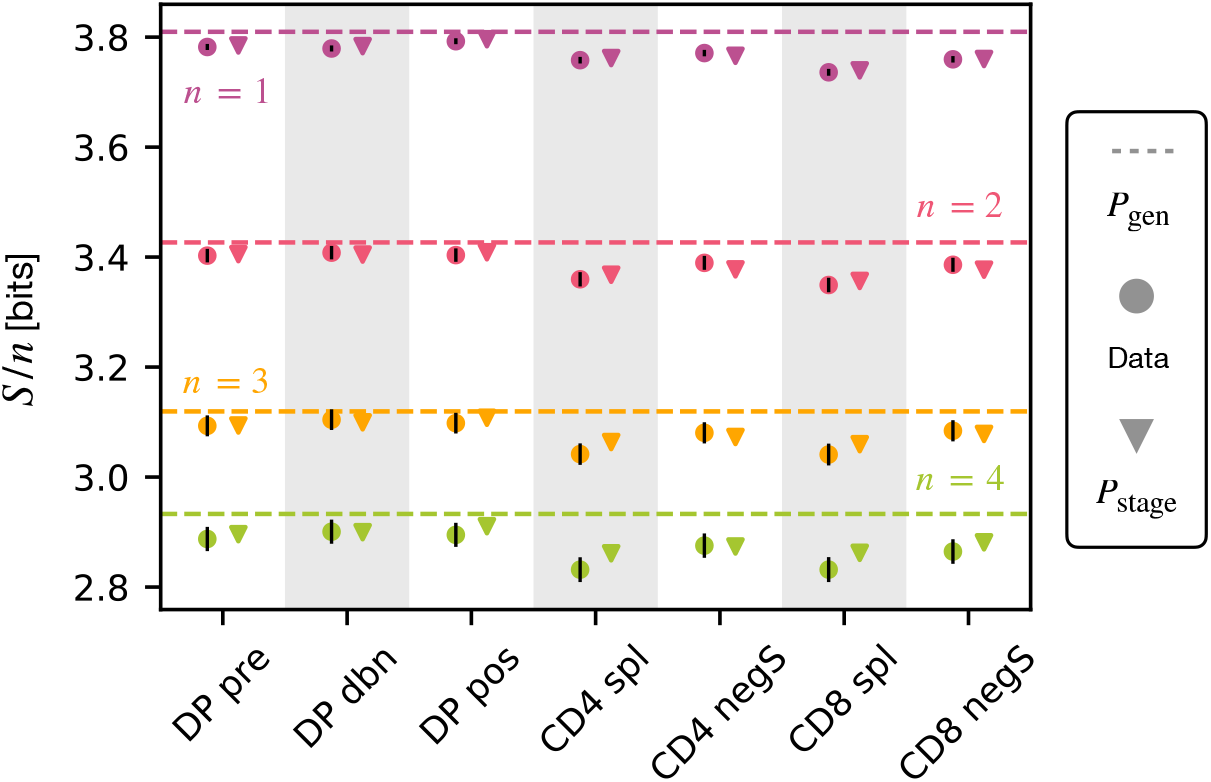
NSB estimation of the Shannon entropy *S* normalized by *n*, associated to *n*-gram distributions within the CDR3 *β* chain of unique clonotypes from the different maturation stages.

**FIG. S10:**
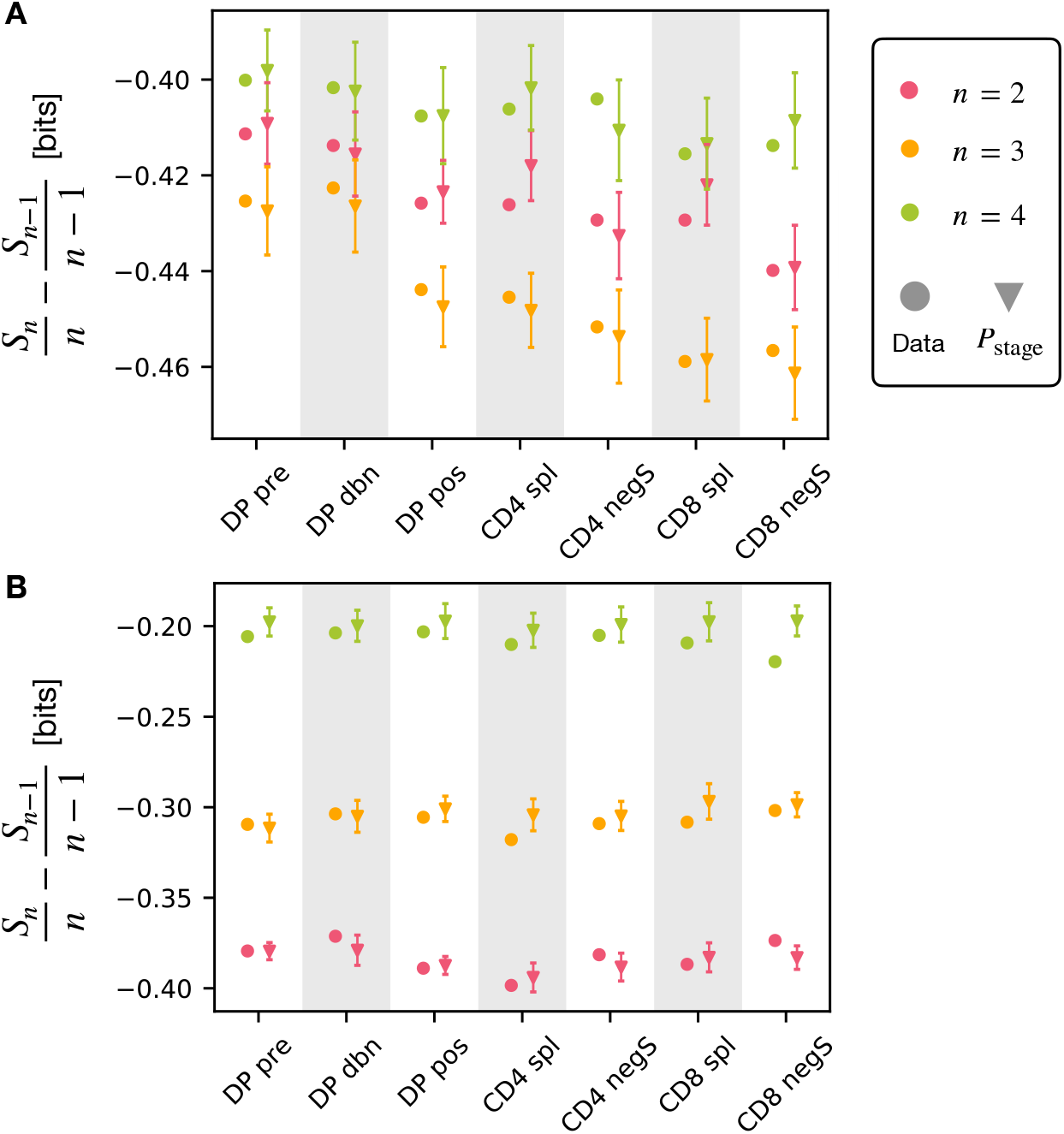
Decrease in entropy per symbol between *n* and *n* – 1 grams. **A** For the *α* chain the decreases are comparable. **B** For the *β* chain the decreases get smaller with *n*.

**FIG. S11:**
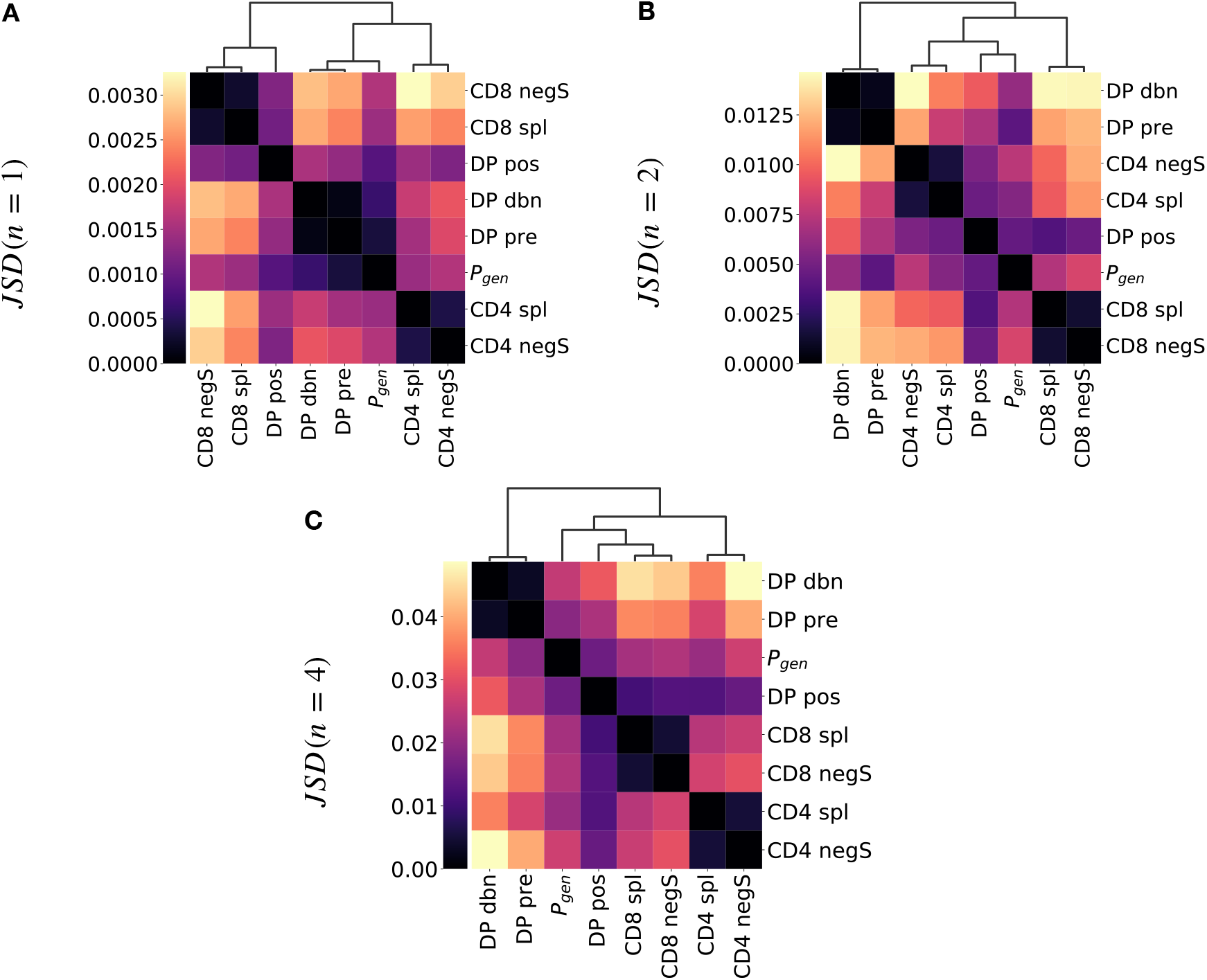
Jensen-Shannon divergence (Eq. 6) for different *n*-gram distributions estimated on *α* chain CDR3 synthetic repertoires for different maturation stages. The dendrogram is computed with the Ward method. **A** *n* = 1. **B** *n* = 2. **C** *n* = 4.

**FIG. S12:**
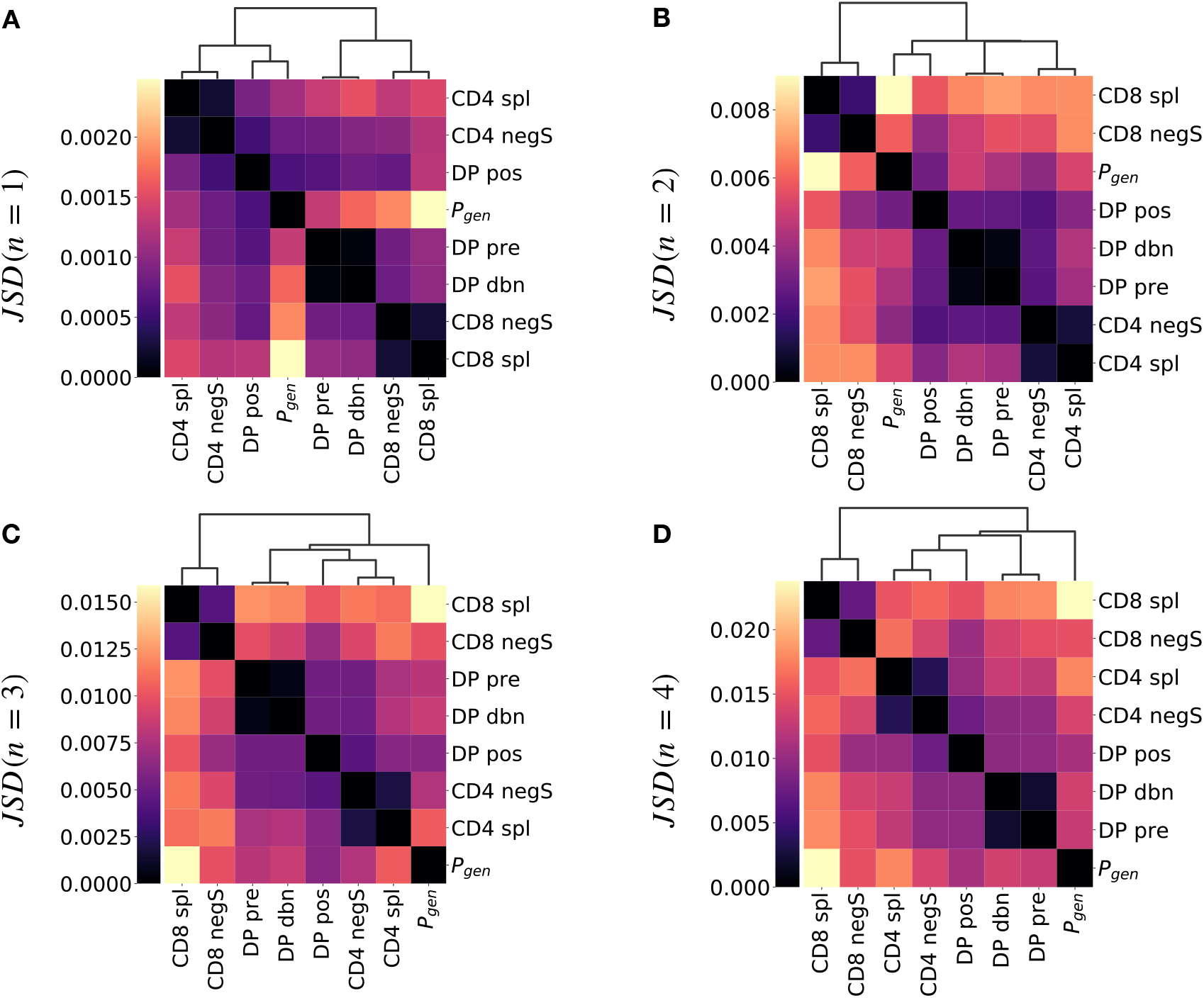
Jensen-Shannon divergence (Eq. 6) for different *n*-gram distributions estimated on *β* chain CDR3 synthetic repertoires for different maturation stages. **A** *n* = 1. **B** *n* = 2. **C** *n* = 3. **D** *n* = 4.

**FIG. S13:**
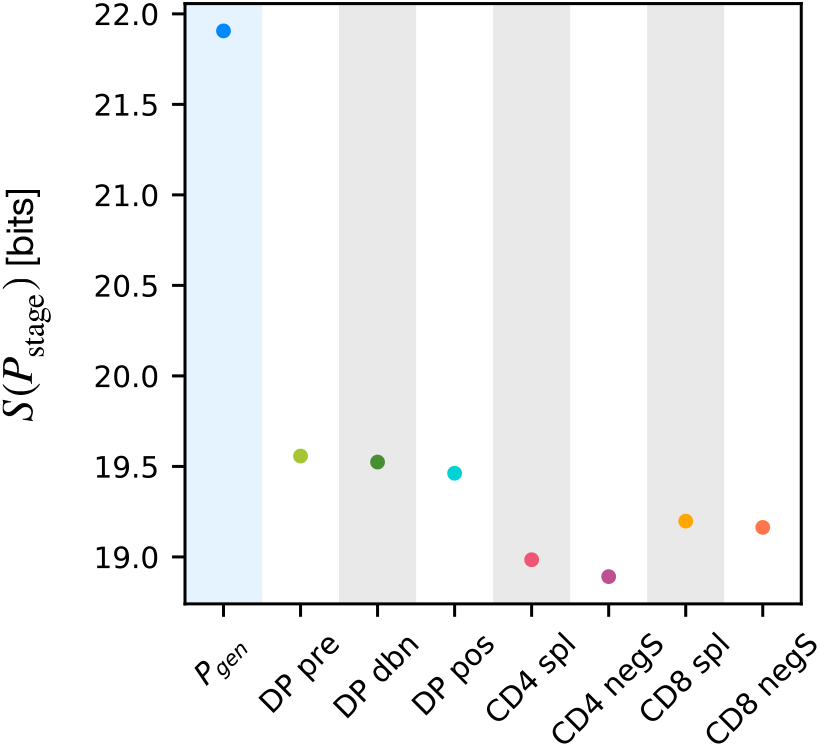
Shannon entropy estimation associated to the full *P*_stage_ model for the *α* chain (Eqn. 5) for the different selection models and the *P*_gen_ generation model.

**FIG. S14:**
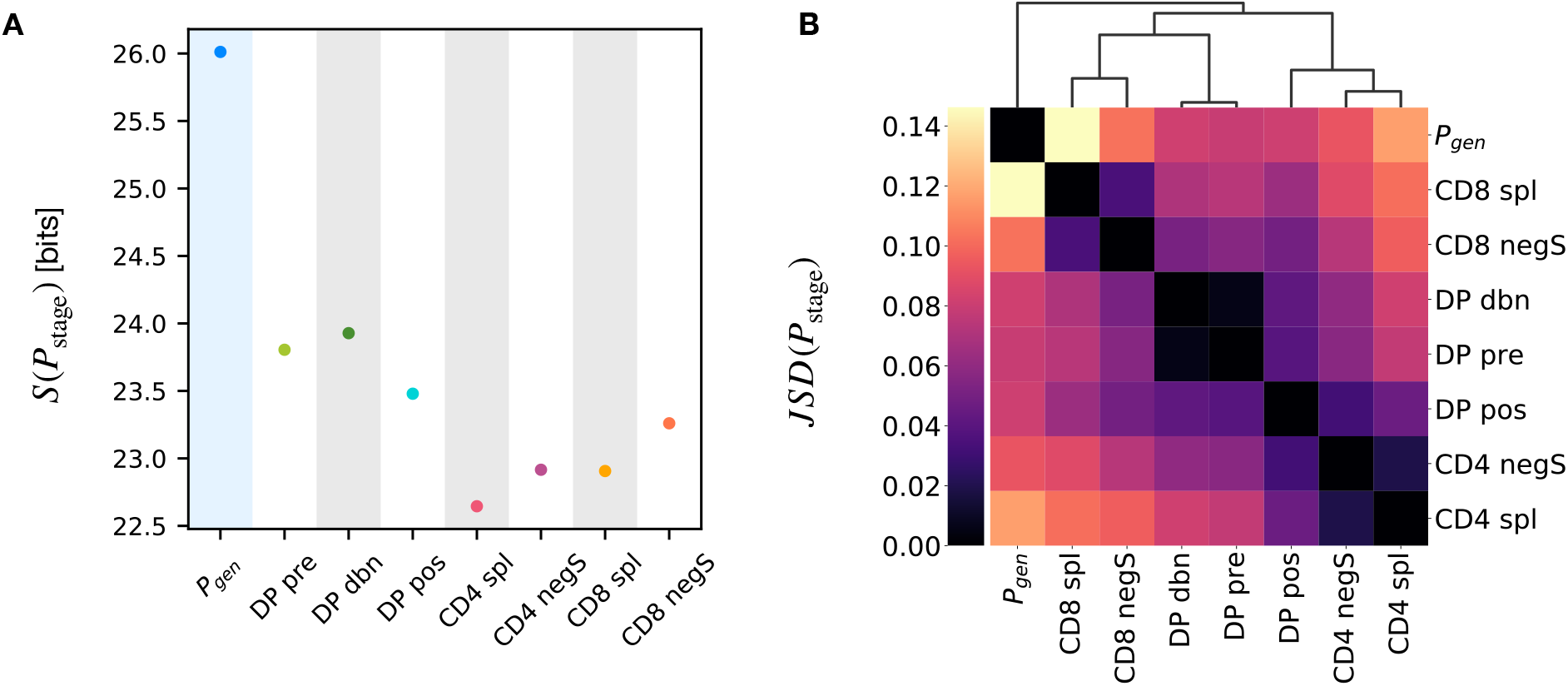
**A.** Shannon entropy estimation associated to the full *P*_stage_ model for the *β* chain (Eqn. 5) for the different selection models and the *P*_gen_ generation model. **B.** Jensen-Shannon divergence for the *β* chain *P*_stage_ models.

**FIG. S15:**
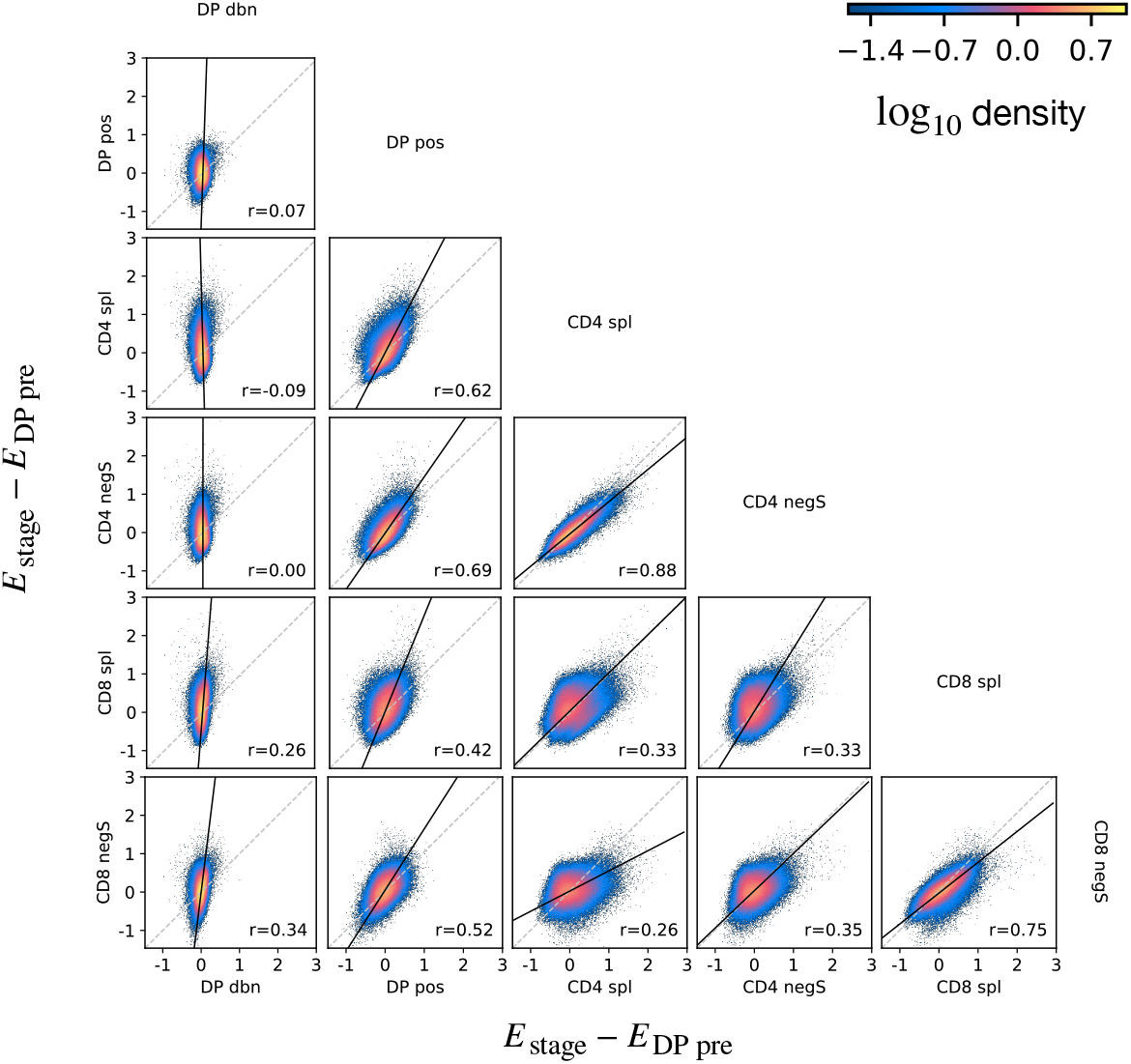
Density scatterplots of the energy differences for *β* chain selection models at different maturation stages and *DP pre* selection models. All selection models are calculated with respect the same *P*_gen_ distribution.

**FIG. S16:**
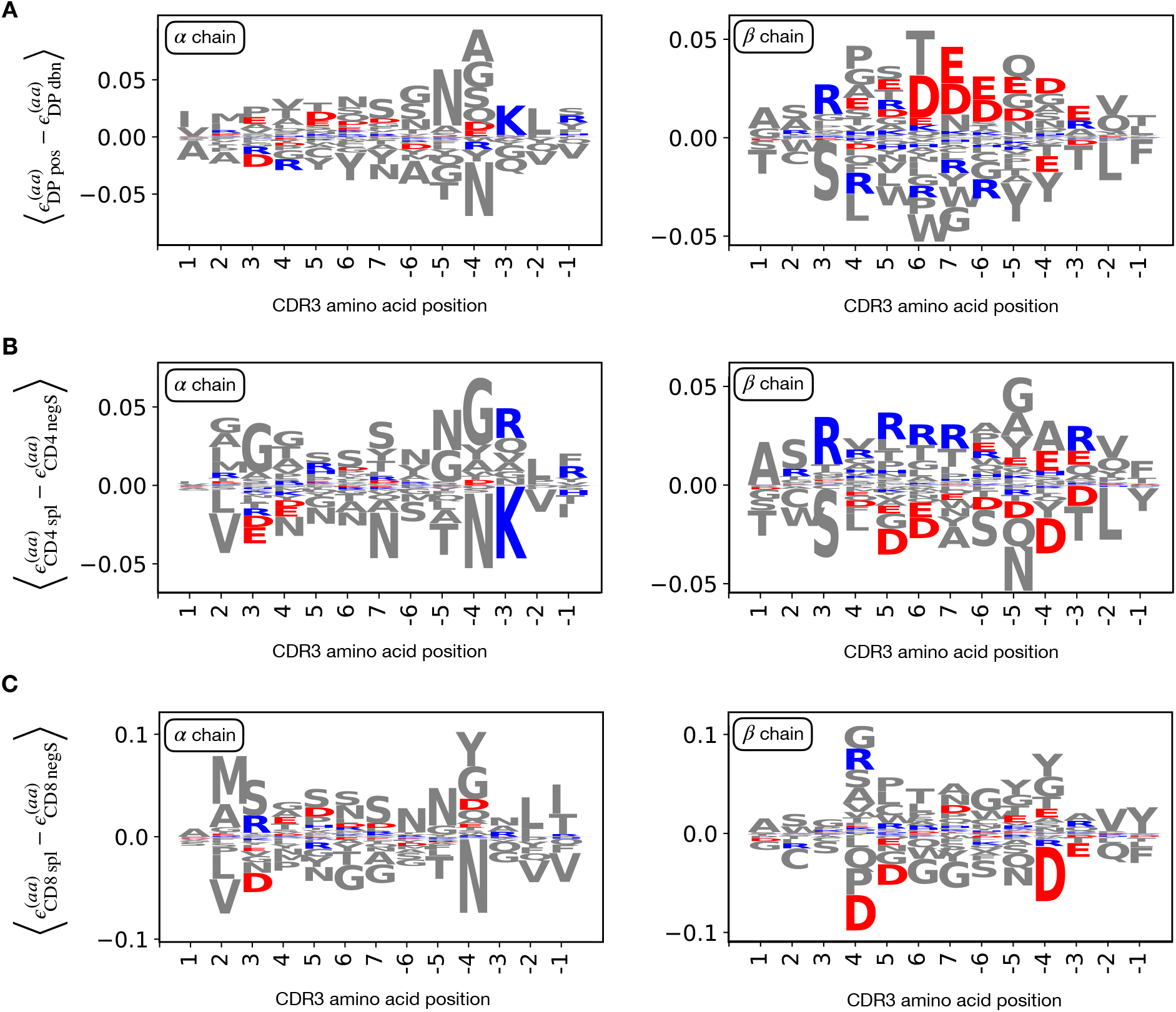
Logo plots for the CDR3 amino acid usage inferred by the model, from the left (positive position indexes) and from the right (negative), omitting the first and the last one. Here the quantity 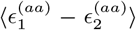 represents the average difference between weights associated to amino acid *aa* at the given position by *P*_stage_ models (see Materials and Methods). Analogously with the energy difference, a negative difference implies the feature is favoured in stage 1, vice versa for stage 2. We follow the color scheme from [49] to highlight the charge properties (red for positive charge, blue for negative charge). **A** Weights difference between stages DP pos and DP dbn. For the *β* chain we see a reduction of positively charged amino acid in DP pos. **B** Stages CD4 spl and CD4 negS. Conversely, here the CD4 spl stage show enhancement in positively charged for the *β* chain. **C** Stages CD8 spl and CD8 negS. We observe just a slight enhancement of positively charged amino acid in CD8 spl.

**FIG. S17:**
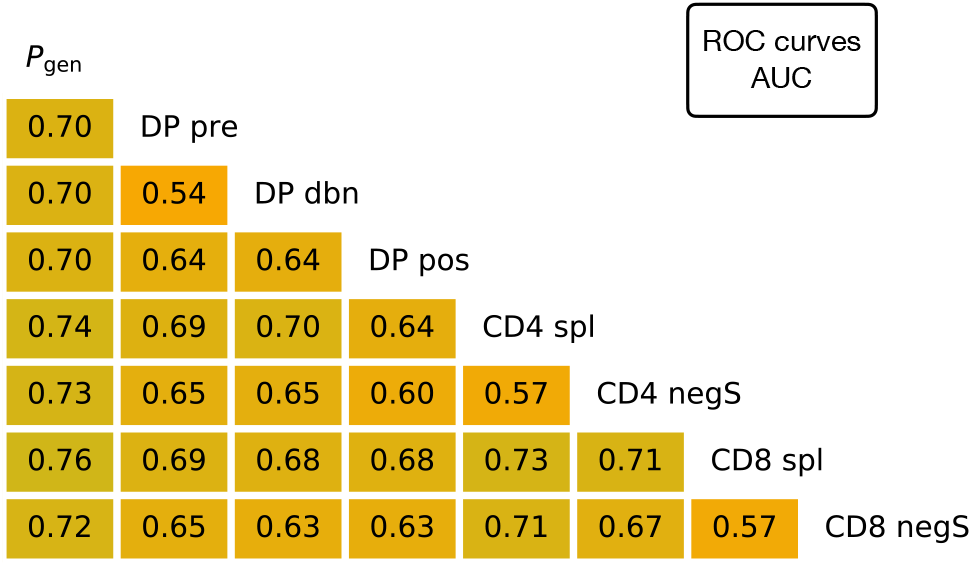
AUC values computed from the ROC curves of the linear classifiers for *β* TCR sequences between pairs of maturation stages. The training/testing set is a random subsample containing 70%/30% of the full dataset at a given maturation stage.

